# LRRK2 regulates ArfGAP1 membrane localization, activity and neuronal toxicity via phosphorylation within its lipid-sensing ALPS2 motif

**DOI:** 10.64898/2026.01.12.699049

**Authors:** Md. Shariful Islam, Valentin Cóppola-Segovia, Alessandra Musso, Darren J. Moore

**Affiliations:** Department of Neurodegenerative Science, Van Andel Institute, Grand Rapids, Michigan 49503, United States; Laboratory of Molecular Neurodegenerative Research, Brain Mind Institute, Swiss Federal Institute of Technology (EPFL), 1015 Lausanne, Switzerland

## Abstract

Mutations in the *leucine-rich repeat kinase 2* (*LRRK2*) gene cause late-onset, autosomal dominant Parkinson’s disease (PD). *LRRK2* encodes a multi-domain protein containing a Roc GTPase domain and a serine/threonine-directed protein kinase domain, with PD-linked mutations known to enhance LRRK2 kinase activity and neuronal toxicity. Our previous studies identified the Golgi protein, ADP-Ribosylation Factor GTPase-Activating Protein 1 (ArfGAP1), as a novel modifier of LRRK2-induced cellular toxicity, where it can serve as a GAP-like protein and a robust kinase substrate of LRRK2. Here, we further explore the phosphorylation of ArfGAP1 by LRRK2 and its functional consequences. LRRK2 mediates the robust phosphorylation of ArfGAP1 within its lipid-sensing ALPS2 motif at residues Ser284, Thr291 and Thr292. We mutated these three phosphorylation sites, either alone or combined, to create hydrophobic phospho-null or charged phospho-mimicking versions of ArfGAP1. We find that modulating ArfGAP1 phosphorylation impairs its normal capacity to induce Golgi fragmentation upon overexpression in neural cells. Blocking phosphorylation impairs ArfGAP1-induced neurite outgrowth inhibition in primary neurons and protects against the neurotoxic effects of PD-linked G2019S LRRK2. ArfGAP1 interactome analysis in neural cells identifies 114 putative interacting proteins with a proportion of these unexpectedly localized to mitochondria, including the outer membrane proteins Voltage-Dependent Anion Channel (VDAC) 1-3. An ArfGAP1 triple phospho-mimic displays an increased interaction with mitochondrial VDACs owing to the redistribution of ArfGAP1 from the *cis*-Golgi to the cytoplasm. Promoting ArfGAP1 phosphorylation also blocks the formation of Golgi-derived vesicles following mild ER stress. Our data provides evidence for a complex functional interaction between LRRK2 and ArfGAP1 that serves to regulate ArfGAP1 subcellular localization, protein interactions, activity and neuronal toxicity via LRRK2-mediated phosphorylation of its membrane-binding ALPS2 motif. Our findings support additional validation of ArfGAP1 as a putative therapeutic target for modulating *LRRK2*-linked PD.

## Introduction

Parkinson’s disease (PD) is a common and progressive neurodegenerative movement disorder that is thought to result from a complex interplay between aging, genetic factors and environmental exposure [1, 2]. While PD typically occurs as a sporadic disease, 5-10% of cases occur in a familial manner with mutations in at least 20 genes identified to cause monogenic forms of PD [3]. Among these familial cases, mutations in the *leucine-rich repeat kinase 2* (*LRRK2*) gene cause late-onset, autosomal dominant PD [4, 5]. In addition, genome-wide association studies implicate common non-coding variation at the *LRRK2* locus in sporadic PD risk [6, 7], indicating that *LRRK2* represents a pleomorphic risk gene for PD. At least seven mutations in *LRRK2* are known to be pathogenic based upon clear segregation with disease in *LRRK2*-linked families, including N1437H, R1441C/G/H, Y1699C, G2019S and I2020T, with G2019S representing the most frequent mutation [8–12]. LRRK2 therefore represents an important and promising therapeutic target for familial and sporadic PD.

LRRK2 belongs to the ROCO protein family and contains multiple domains including a central catalytic region consisting of Ras-of-Complex (Roc) GTPase and serine/threonine-directed protein kinase domains separated by a C-terminal-of-Roc (COR) linker region [12, 13]. LRRK2 can function as a kinase and GTPase, with the capacity for GTP-binding serving to regulate kinase activity [12]. A number of substrates have been identified for LRRK2 kinase activity, with a subset of ∼14 Rab GTPases (i.e., Rab8A/10/12/29) being the best characterized to date especially within mammalian cells [13–15]. Additional LRRK2 kinase substrates have also been identified mostly using *in vitro* assays, including ArfGAP1 [16, 17], RPS15 [18], β-tubulin [19, 20], MARK1 [20], NSF [21], snapin [22], synaptojanin-1 [23], auxilin 1 [24], RGS2 [25], FoxO1 [26] and Futsch [23, 27]. Familial PD mutations in LRRK2 tend to cluster within the Roc, COR and kinase domains. Notably, familial PD mutations share the capacity to enhance kinase activity, either directly via the kinase activation loop (G2019S, I2020T) or indirectly via the Roc-COR tandem domain (R1441C/G/H, Y1699C) presumably by impairing GTP hydrolysis and prolonging the GTP-bound state [12, 13]. Familial PD mutations also commonly inhibit neurite outgrowth and induce cell death in primary neuronal culture models via a kinase-dependent mechanism [12, 28–30]. The GTPase and kinase activity of LRRK2 are both important for the development of PD and further understanding the accessory factors and substrates that regulate these activities will provide key insight for developing therapeutic strategies to attenuate LRRK2 activity in PD.

Our previous studies have identified ADP-ribosylation factor GTPase-activating protein 1 (ArfGAP1) as a novel modifier of LRRK2 activity and cellular toxicity. The deletion of *GCS1*, an ortholog of mammalian ArfGAP1, was originally identified as a suppressor of human LRRK2-induced cellular toxicity in yeast [31]. Subsequent studies revealed that ArfGAP1 gene silencing rescues the inhibition of neurite outgrowth induced by G2019S LRRK2 in rodent primary cortical neurons, whereas co-expression of wild-type LRRK2 and ArfGAP1 synergistically promotes neurite outgrowth deficits [16, 17]. ArfGAP1 expression alone can also reduce neurite length in a manner that depends on endogenous LRRK2 [16]. These data indicate the complex functional relationship between ArfGAP1 and LRRK2 and suggest that ArfGAP1 is critically required for mediating the pathogenic actions of mutant LRRK2. At the molecular level, ArfGAP1 physically interacts with LRRK2 and promotes its GTP hydrolysis activity and unexpectedly enhances LRRK2 kinase activity [16, 17]. ArfGAP1 also serves as a robust and direct substrate of LRRK2-mediated phosphorylation [16, 17]. Therefore, ArfGAP1 represents an intriguing accessory protein for LRRK2 that may function as a GAP-like protein to modulate LRRK2 activity and as a kinase substrate of LRRK2. We hypothesize that ArfGAP1 could mediate LRRK2-induced toxicity either by i) serving upstream as a GAP-like protein to enhance the GTPase activity of LRRK2, or ii) serving downstream as a substrate of LRRK2-dependent phosphorylation. We have elected to explore the role of ArfGAP1 phosphorylation as a potential mechanism, since available cryo-EM structural data suggests that dimeric LRRK2 is unlikely to accommodate or require GAPs and guanine nucleotide exchange factors (GEFs) for regulating Roc GTPase activity [32, 33]. As such, LRRK2 has instead been proposed to function as a non-canonical GTPase, specifically via a GTPase-activated-by-dimerization (GAD) mechanism [34, 35].

We and others have shown that LRRK2 can directly phosphorylate ArfGAP1 *in vitro* [16, 17], but the impact of phosphorylation on ArfGAP1 cellular function remains to be determined. ArfGAP1 is known to function as a GAP for the Golgi-localized small GTPase Arf1, that mediates the dissociation of the COPI coatomer complex from Golgi-derived vesicles, a prerequisite for vesicle fusion with target compartments such as the endoplasmic reticulum (ER) [36–40]. As such, overexpression of ArfGAP1 in mammalian cells induces the redistribution of the entire Golgi complex to the ER via vesicular intermediates [39, 41, 42]. As ArfGAP1 is required for LRRK2-induced neuronal toxicity, and ArfGAP1 silencing in neurons is generally well-tolerated [16, 17], ArfGAP1 inhibition may offer an alternative therapeutic target for attenuating LRRK2 activity and toxicity in PD. Here, we identify LRRK2-specific phosphorylation sites in ArfGAP1 and explore the effects of modulating these phosphorylation sites on its subcellular localization, Golgi morphology and sorting, neurite outgrowth and protein interactions. Our data further elucidates the functional interaction between LRRK2 and ArfGAP1, and the significance of ArfGAP1 phosphorylation.

## Results

### LRRK2 phosphorylates ArfGAP1 at Ser284 and Thr291 or Thr292

To explore the functional interaction between LRRK2 and ArfGAP1, we focused on defining the impact of LRRK2-mediated ArfGAP1 phosphorylation. We have previously demonstrated that ArfGAP1 serves as a robust substrate of LRRK2-mediated phosphorylation by *in vitro* radioactive kinase assay [16]. However, the identification and quantitation of LRRK2-specific ArfGAP1 phosphorylation sites was lacking in our previous work. A prior study identified six putative phosphorylation sites (S155, T189, T216, S246, S284, T292) within ArfGAP1 by mass spectrometry yet no quantitative analysis was provided, and it was unclear which LRRK2 variants were used in these assays, including whether comparison to a kinase-inactive LRRK2 negative control was included [17]. Furthermore, the simultaneous mutation of all six phospho-sites to alanine was required to inhibit LRRK2-mediated ArfGAP1 phosphorylation, suggesting that none of these six residues serve as a dominant phospho-site [17]. To initially locate the most abundant phosphorylation sites within ArfGAP1, we conducted *in vitro* radioactive kinase assays using [^33^P]-γ-ATP together with immunopurified recombinant full-length FLAG-tagged human LRRK2 (WT, G2019S or D1994A) and a series of GST-tagged ArfGAP1 protein fragments (residues 1-136, 137-415, 137-251, 252-359 and 360-415) (**Fig. 1A**). We initially purified each GST-tagged ArfGAP1 protein from *E.coli* using glutathione-sepharose affinity columns and confirmed their relative purity by SDS-PAGE and Coomassie staining (**Fig. 1B**) before subjecting each protein fragment to *in vitro* kinase assays with LRRK2 (**Fig. 1C-G**). We find that ArfGAP1 proteins containing residues 137-415 and 252-359 are robustly phosphorylated by G2019S LRRK2 and to a lesser extent by WT LRRK2, relative to kinase-inactive D1994A LRRK2 or in the absence of LRRK2 (**Fig. 1C, G**). Other ArfGAP1 proteins (residues 1-136, 137-251 and 360-415) are only modestly phosphorylated by G2019S LRRK2 relative to other LRRK2 variants (**Fig. 1D-F**). Importantly, the N-terminal catalytic GTPase-activating domain of ArfGAP1 (contained within residues 1-136), serves as a poor substrate of LRRK2 (**Fig. 1E**). These data suggest that the majority of LRRK2-mediated phosphorylation signal minimally resides within residues 252-359 of ArfGAP1 (GAP-F5 fragment), although additional less abundant sites are likely to exist outside of this region as suggested by higher phosphorylation of the GAP-F1 fragment (137-415; **Fig. 1C**).

**Figure 1.**
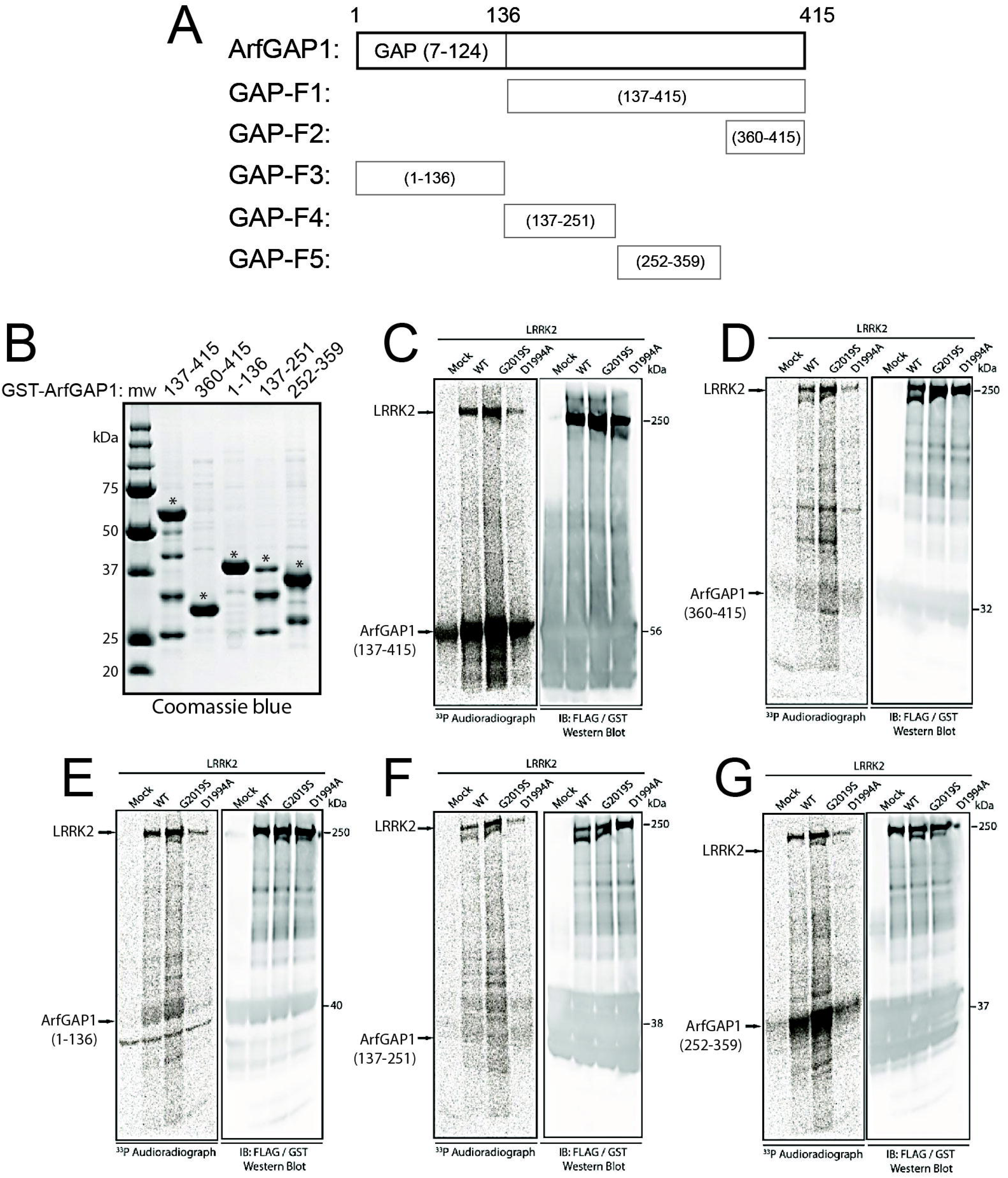
LRRK2-mediated phosphorylation of isolated ArfGAP1 domains. Schematic diagram of GST-tagged ArfGAP1 domain constructs (F1-F5, **A**) and their purity/levels on SDS-PAGE gels with Coomassie staining following purification using Glutathione-sepharose pull-down (**B**). **C-G**) Phosphorylation of each recombinant ArfGAP1 domain fragment by IP full-length human FLAG-tagged LRRK2 (WT, G2019S, D1994A) by *in vitro* kinase assay with [^33^P]-γ-ATP. Phosphorylation analysis of ArfGAP1 domains reveals that fragments 137-415 (F1, **C**) and 252-359 (F5, **G**) are abundantly phosphorylated by WT and G2019S LRRK2, with minimal phosphorylation of fragments 360-415 (F2, **D**), 1-136 (F3, **E**) and 137-251 (F4, **F**). LRRK2 autophosphorylation is also indicated with D1994A serving as a kinase-inactive control, and a mock FLAG IP (no LRRK2) serving as a control for non-specific incorporation of ^33^P. Indicated are representative ^33^P autoradiographs (left panel) and corresponding Western blots (right panel) co-probed with anti-GST (ArfGAP1) and anti-FLAG (LRRK2) antibodies to indicate equivalent protein loading. Molecular mass is indicated in kilodaltons (kDa). Positions of full-length LRRK2 and ArfGAP1 domains are indicated with arrows.

To identify the major phosphorylation sites in full-length ArfGAP1, we conducted similar *in vitro* kinase assays with non-radioactive ATP and subjected samples to mass spectrometry for quantitative analysis of ArfGAP1 phospho-peptides, comparing the effects of WT, G2019S and D1994A LRRK2. *In vitro* kinase assays with recombinant GST-tagged human LRRK2 variants (ΔN-LRRK2, residues 970-2527) and full-length rat ArfGAP1 were subjected to in-solution digestion to generate peptide mixtures for nano-LC-MS/MS analysis (Q Exactive, Thermo Fisher) using a 2-hour gradient time, as described [43]. Mass spectrometry analysis of *in vitro* phosphorylated rat ArfGAP1 identifies four major sites (pT145, pT189, pT230, pT292) that are modified by LRRK2 to varying degrees (**Fig. 2A**). We confirm the ∼2-fold enhanced phosphorylation of ArfGAP1 at pT145, pT230 and pT292 by G2019S LRRK2 relative to WT LRRK2, whereas pT189 levels are similar between WT and G2019S LRRK2 (**Fig. 2B**). Importantly, kinase-inactive D1994A LRRK2 fails to detectably phosphorylate these four sites within ArfGAP1 (**Fig. 2B**). Notably, we detect two threonine phosphorylation sites (pT189, pT292) identified in a prior study [17], and two novel threonine sites (pT145, pT230), but we fail to detect four previously identified LRRK2-specific phosphorylation sites (pS155, pT216, pS246, pS284) in these assays [17]. Of note, rat ArfGAP1 contains a N155 residue instead of S155 present in the human and mouse protein. MS/MS spectra are shown for the most abundant ArfGAP1 phospho-peptide (^289^DVT[**pT**]FFSGK^297^) that supports phosphorylation at T292 in the presence of WT and G2019S LRRK2 (**Fig. 2C**). Interestingly, of the combined phospho-sites identified by us and others, S155, T189, T230, S246 and S284 are known phosphorylation sites in mammalian ArfGAP1 according to the *PhosphoSitePlus^®^* database, whereas intriguingly T291 rather than T292 is also a known phosphorylation site. Importantly, of the phospho-sites identified by MS, pT292 lies within the GAP-F5 fragment (residues 252-359) of ArfGAP1 that is robustly phosphorylated by LRRK2 (**Fig. 1G**). We selected two additional putative ArfGAP1 phosphorylation sites (S284, T291) for further analysis, since pS284 was identified by a prior study [17] and also lies within the GAP-F5 fragment, whereas the known phospho-site T291 is adjacent to T292 that makes it challenging to assign precise phospho-site localization within a single phospho-peptide.

**Figure 2.**
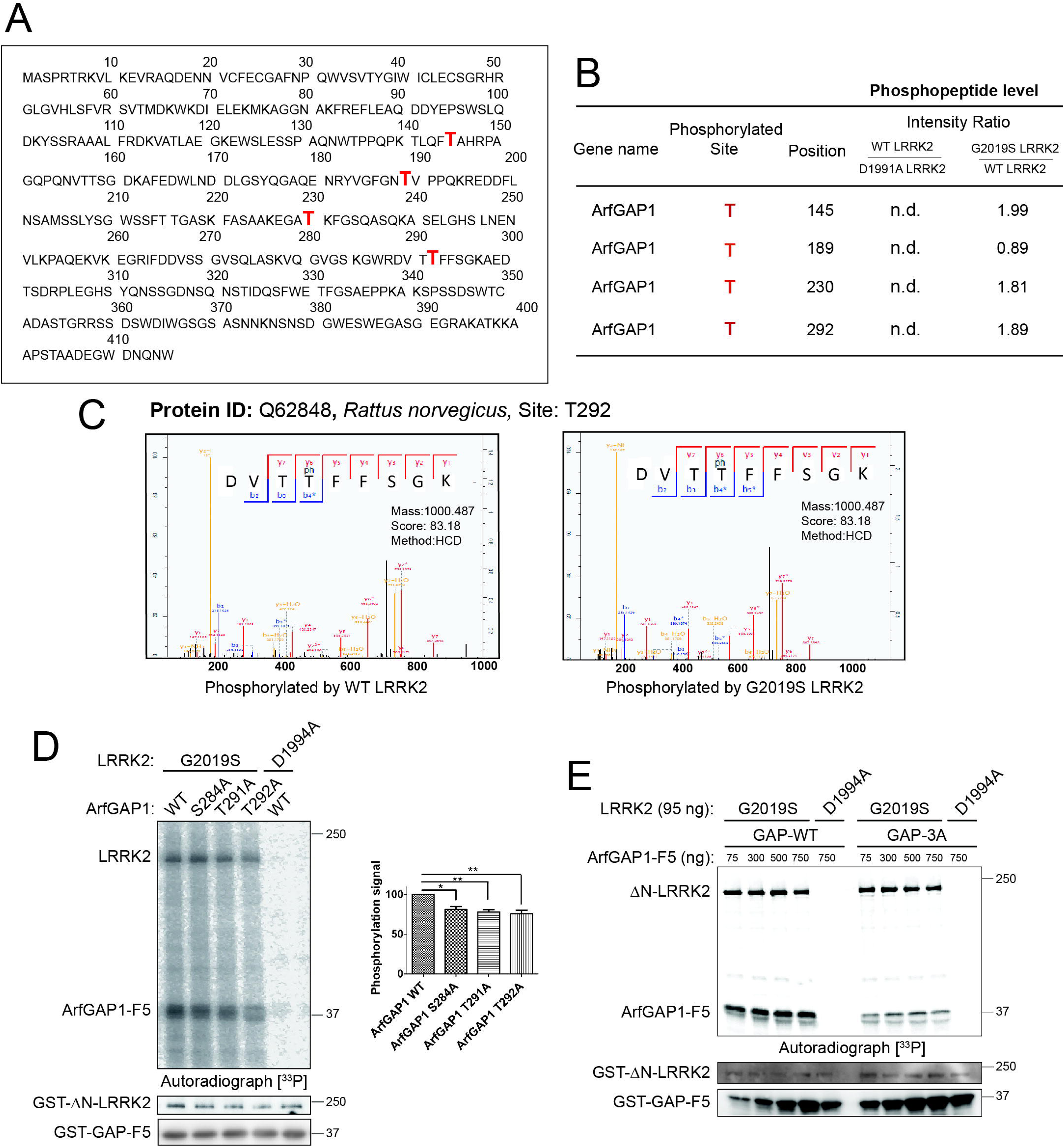
Identification of LRRK2-mediated ArfGAP1 phosphorylation sites. **A**) Full-length protein sequence of rat ArfGAP1 with phosphorylated sites indicated in red. Mass spectrometry-based phospho-proteomic analysis identifies four major sites (T145, T189, T230, T292) that are phosphorylated by WT and G2019S LRRK2. **B**) Table of significantly regulated ArfGAP1 phosphopeptides mediated by WT LRRK2 compared to kinase-inactive D1994A or G2019S LRRK2. pT145, pT230 and pT292 are increased ≥1.8-fold by G2019S relative to WT LRRK2. **C**) Selected MS/MS spectra for ArfGAP1 phosphopeptide (DVTp**T**FFSGK; pT292) for WT or G2019S LRRK2. The respective phosphorylated site, the corresponding Uniprot ID and the Andromeda score are shown. **D**) *In vitro* phosphorylation of GST-tagged ArfGAP1 domain (F5, residues 252-359) WT or single phospho-null mutants (S284A, T291A, T292A) by recombinant GST-tagged human LRRK2 (ΔN, residues 970-2527, D1994A or G2019S) with [^33^P]-γ-ATP. Representative ^33^P autoradiograph indicating ArfGAP1 (and LRRK2) phosphorylation signal is shown, and Western blots probed with anti-GST antibody. Graph indicates densitometric analysis of each ArfGAP1-F5 phosphorylation signal (with G2019S LRRK2) normalized to total ArfGAP1-F5 levels, with bars representing the mean ± SEM (*n* = 3 experiments). \**P*<0.05 or \*\**P*<0.01 compared to WT ArfGAP1 signal by one-way ANOVA with Dunnett’s multiple comparisons test, as indicated. **E**) *In vitro* phosphorylation of increasing quantities of GST-tagged ArfGAP1-F5 domain WT or triple phospho-null mutant (3A: S284A/T291A/T292A; 75-750 ng) by GST-tagged LRRK2 (D1994A or G2019S). Representative ^33^P autoradiograph and Western blots probed with anti-GST antibody are shown. Molecular mass is indicated in kilodaltons (kDa). Positions of ΔN-LRRK2 and ArfGAP1-F5 domain are indicated.

To distinguish which of these three sites are the dominant phosphorylation site of ArfGAP1, we generated recombinant GST-tagged ArfGAP1 proteins (GAP-F5 fragment) harboring individual phospho-null alanine mutations (S284A, T291A and T292A) for *in vitro* radioactive kinase assays with recombinant GST-tagged ΔN-LRRK2 (970-2527). We find that each phospho-null mutation within the GAP-F5 protein significantly yet partially reduces G2019S LRRK2-mediated phosphorylation relative to WT GAP-F5, however, none of these three residues serve as a dominant phospho-site as they have equivalent effects (**Fig. 2D**). D1994A LRRK2 fails to appreciably phosphorylate the WT GAP-F5 protein in this assay (**Fig. 2D**). These data suggest that multiple sites within ArfGAP1, and particularly residues 252-359, are simultaneously phosphorylated by LRRK2. To further confirm these three ArfGAP1 phospho-sites, we generated a recombinant GST-tagged GAP-F5 protein containing all three phospho-null alanine mutations (referred to as 3A: S284A/T291A/T292A). The 3A mutation in GAP-F5 protein effectively blocks the increase in G2019S LRRK2-mediated phosphorylation in response to increasing amounts of GAP-F5 protein substrate, compared to the concentration-dependent increase in WT GAP-F5 phosphorylation, by *in vitro* radioactive kinase assay (**Fig. 2E**). D1994A LRRK2 fails to phosphorylate the WT or 3A GAP-F5 proteins in these assays (**Fig. 2E**), confirming the specificity of LRRK2 kinase activity for ArfGAP1. It is not possible to reliably determine whether T291 or T292 are phosphorylated by LRRK2 as alanine mutations of one site most likely impact the phosphorylation motif of the other site, whereas definitively resolving phosphorylation sites between adjacent identical residues (i.e., pTT or TpT) in a single phospho-peptide by mass spectrometry is not possible. In subsequent experiments, we have elected to mutate both T291 and T292 together to overcome this potential issue. Our attempts to verify these phospho-sites in cells co-expressing ArfGAP1 with LRRK2 variants combined with mass spectrometry analysis of immunoprecipitated ArfGAP1 were unsuccessful, most likely due to the low abundance and/or compartment-specific localization of these LRRK2-specific phosphorylation sites for detection in whole cell extracts. Collectively, our data suggests that S284 and T291 or T292 represent major sites within ArfGAP1 that are phosphorylated by LRRK2.

Interestingly, the S284 and T291/T292 phospho-sites are localized within a unique motif termed ArfGAP1 lipid-packing sensor 2 (ALPS2), that consists of residues 264-295 [44, 45]. ArfGAP1 also contains a similar ALPS1 motif (residues 199-234), with both ALPS motifs localized to the non-catalytic C-terminal portion of ArfGAP1 relative to the GAP catalytic domain localized within the first 136 residues [44, 45]. The ALPS 1 and 2 motifs bind preferentially to highly curved membranes and couple Arf1 GTP hydrolysis to COPI coat-induced membrane curvature [44, 45]. The ALPS2 motif is normally unstructured but forms an amphipathic α-helix at the surface of highly curved membranes, with the α-helix containing an abundance of serine and threonine residues in its polar face [45]. These Ser/Thr residues are critical for the sensitivity of the ALPS2 motif to membrane curvature [45]. Notably, S284, T291 and T292 are incorporated within this α-helix of ALPS2, with the equivalent positions of S284 and T291 also being highly conserved in the ALPS1 motif [45]. The pT230 site identified by mass spectrometry above also represents a highly conserved residue within ALPS1 [45]. It is possible that LRRK2-dependent ArfGAP1 phosphorylation serves to regulate the sensitivity of the ALPS2 motif for binding with highly curved membranes, such as Golgi-derived vesicles, and therefore may modulate Arf1-dependent Golgi-to-ER retrograde transport.

### Impact of LRRK2-specific ArfGAP1 phosphorylation sites on Golgi complex integrity

ArfGAP1 is known to promote the GTP hydrolysis activity of Arf1 that is required for dissociation of the COPI coatmer complex from Golgi-derived vesicles, a prerequisite for vesicle fusion with target compartments such as the ER [37–39]. The overexpression of wild-type ArfGAP1 in mammalian cells leads to excessive Arf1 GTP hydrolysis that can impair Golgi complex integrity due to the increased redistribution of Golgi-derived vesicles to the ER [41, 42]. To explore the impact of ArfGAP1 phosphorylation on this Arf1-dependent activity, we evaluated Golgi morphology in human SH-SY5Y neural cells overexpressing ArfGAP1 phospho-site variants by substituting either uncharged hydrophobic phospho-null (Alanine: A) or negatively-charged phospho-mimicking (Aspartic Acid: D) residues at Ser284, T291 and T292. Golgi morphology therefore serves as a useful functional readout of ArfGAP1 activity. Fluorescence microscopy reveals the effects of ArfGAP1 variants on Golgi morphology using the *cis*-Golgi marker GM130, resulting in either a normal intact perinuclear Golgi complex or more frequently an abnormally fragmented (or absent) Golgi complex (**Fig. 3A**). Quantitative analysis of Golgi fragmentation indicates that overexpression of WT ArfGAP1 induces pronounced Golgi fragmentation in ∼85% of cells, whereas phospho-null (3A) ArfGAP1 significantly attenuates Golgi fragmentation (∼43% cells) (**Fig. 3B**). Unexpectedly, phospho-mimic (3D) ArfGAP1 also similarly reduces Golgi fragmentation (∼55% cells) relative to WT ArfGAP1 (**Fig. 3B**). Similar levels of Golgi complex fragmentation induced by WT ArfGAP1 expression are produced using a range of *cis*- or *trans*-Golgi markers, such as TGN46, Giantin or GOLGA4, similar to GM130 (**Fig. S1**). In addition, the analysis of individual ArfGAP1 phospho-null (S284A, T291A or T292A) or phospho-mimic (S284D, T291D or T292D) mutants alone does not reveal significant differences in Golgi fragmentation relative to WT ArfGAP1 (**Fig. S1**). We further demonstrate that a T291A/T292A double variant of ArfGAP1 induces an intermediate level of Golgi fragmentation relative to WT and 3A ArfGAP1 (**Fig. S1**). Collectively, our data suggests that the combined phosphorylation at S284 and T291 or T292 impairs the activity of ArfGAP1 to induce Arf1-mediated Golgi dispersal, whereas single (S284, T291, T292) mutations have no appreciable effect.

**Figure 3.**
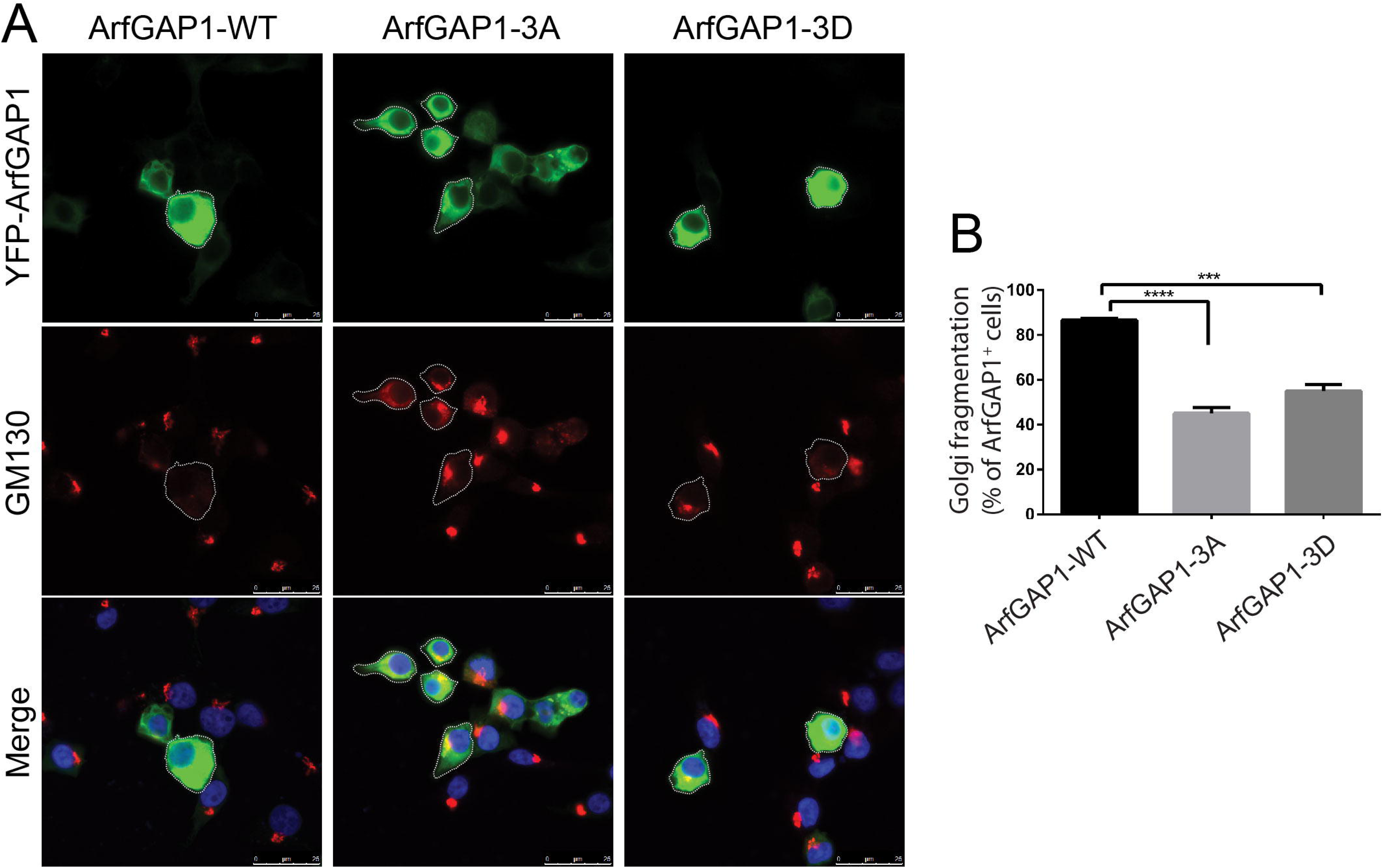
Phosphorylation mutants impair ArfGAP1-induced Golgi fragmentation in neural cells. **A**) SH-SY5Y cells transiently expressing full-length YFP-tagged ArfGAP1 WT, triple phospho-null mutant (3A: S284A/T291A/T292A) or triple phospho-mimic mutant (S284D/T291D/T292D). Fixed cells were subjected to immunofluorescence with an antibody to the *cis*-Golgi marker, GM130. Fluorescence microscopy reveals a robust effect of ArfGAP1 overexpression on Golgi morphology, resulting in a normal (large perinuclear puncta), partially/fully fragmented (dispersed) or absent Golgi complex (white outline). Nuclear DAPI staining is also shown in merged images. **B**) Quantitative analysis of Golgi fragmentation induced by ArfGAP1 overexpression (WT, 3A or 3D). Golgi fragmentation (dispersed or absent) is expressed as a percent of total YFP-ArfGAP1-positive cells for each variant. Bars represent mean ± SEM fragmented Golgi from 120-150 YFP-ArfGAP1-positive cells across at least three independent experiments/cultures. \*\*\**P*<0.001 and \*\*\*\**P*<0.0001 compared to WT ArfGAP1 by one-way ANOVA with Dunnett’s multiple comparisons test, as indicated.

### Impact of LRRK2-specific ArfGAP1 phosphorylation sites on neurite outgrowth

Mutant LRRK2 expression can inhibit neurite outgrowth in cultured primary cortical neurons, particularly axonal processes [16, 46]. We have previously shown that wild-type ArfGAP1 overexpression can similarly inhibit axonal outgrowth and reduce axonal length, whereas ArfGAP1 gene silencing can reverse the effects of G2019S LRRK2 on neurite length [16]. These data indicate that ArfGAP1 plays a role in regulating neurite outgrowth and potentially in neuronal integrity and viability. To evaluate the effects of LRRK2-specific ArfGAP1 phosphorylation sites on neurite outgrowth, we conducted quantitative assays by assessing axonal length in cultured rat primary cortical neurons. Cultures at days-*in-vitro* (DIV) 3 were transiently co-transfected with YFP-tagged ArfGAP1 variants or empty vector and pDsRed-Max-N1 (to mark neuronal processes) at a 10:1 molar ratio and fixed at DIV 6. Fluorescence microscopy was used to measure the longest DsRed-positive neurite (i.e., the axon) from YFP+ (ArfGAP1) or YFP- (empty vector) cortical neurons, as previously described [16]. We find that neurons overexpressing WT ArfGAP1 reveal a marked reduction in neurite length relative to control (empty vector) neurons, whereas unexpectedly, single ArfGAP1 phospho-null mutants (S284A, T291A or T292A) also markedly reduce neurite length to levels similar to WT ArfGAP1 (**Fig. S2**). Furthermore, we find that expression of single ArfGAP1 phospho-mimic mutants (S284D, T291D or T292D) also markedly reduces neurite length similar to WT ArfGAP1 (**Fig. S3**). These neurite length data are consistent with the lack of effect of single phospho-site mutants on ArfGAP1-induced Golgi fragmentation (**Fig. S1**). Accordingly, we compared the effects of triple ArfGAP1 phospho-site mutants (3A or 3D) on neurite length. Remarkably, we find that the 3A mutant partially yet significantly attenuates the impact of ArfGAP1 expression on neurite length relative to the WT or 3D proteins which are similar (**Fig. 4**). Together, these data indicate that blocking the combined phosphorylation of ArfGAP1 at Ser284, T291 and T292 (3A mutant), but not at single sites, impairs neurite outgrowth inhibition consistent with reducing ArfGAP1 activity. Mimicking ArfGAP1 phosphorylation at these sites is sufficient for maximal inhibition of neurite outgrowth, similar to WT ArfGAP1, potentially suggesting that these phosphorylation sites are already saturated in the WT protein in cortical neurons.

**Figure 4.**
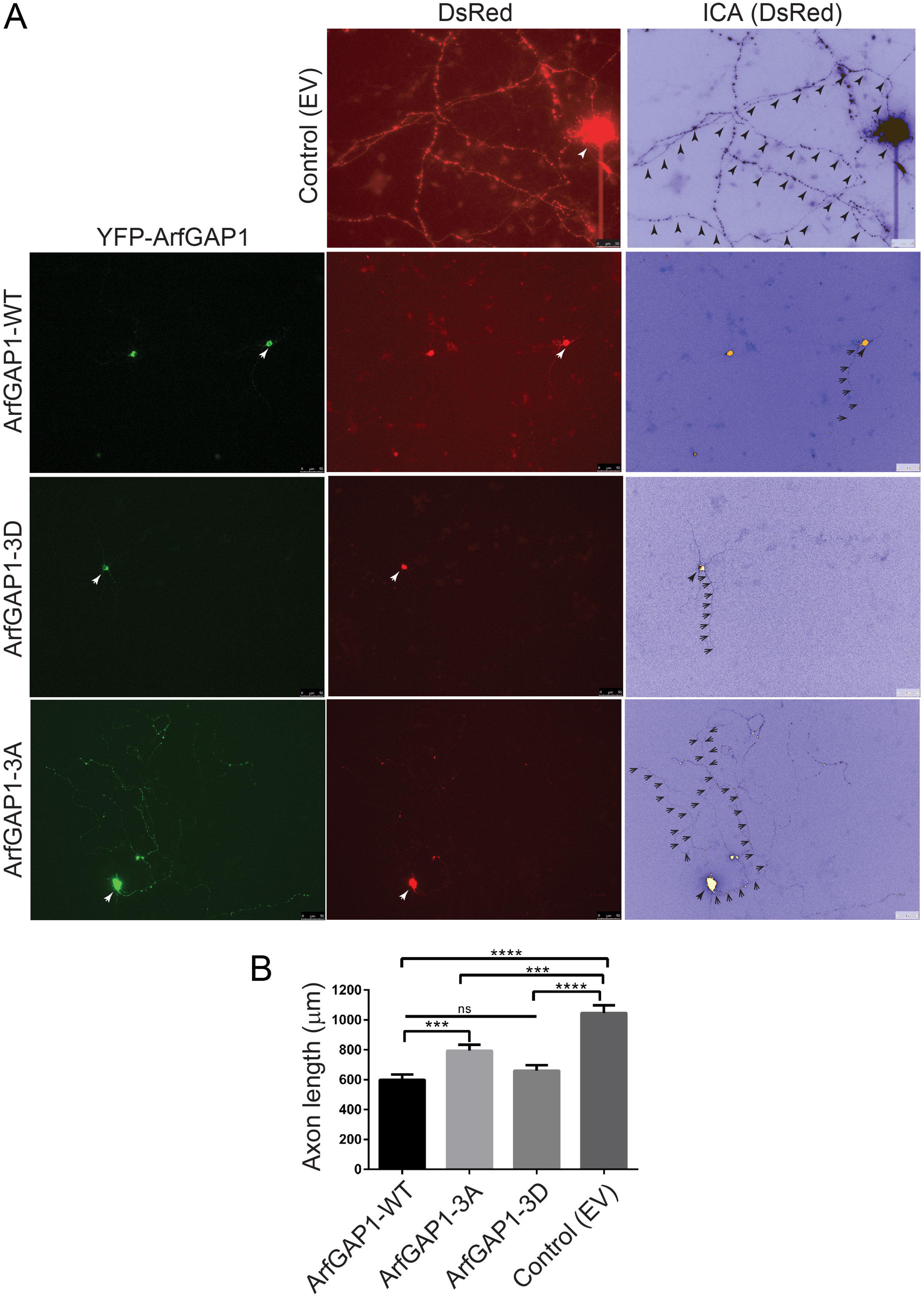
Preventing phosphorylation attenuates the ArfGAP1-induced inhibition of neurite outgrowth in cortical neurons. **A**) Rat primary cortical neurons were co-transfected at DIV3 with YFP-ArfGAP1 (WT, 3A or 3D) or empty vector and DsRed-Max-N1 plasmids, and fixed at DIV6 for confocal fluorescence microscopy analysis. Fluorescent images reveal the co-labeling of cortical neurons with YFP-ArfGAP1 (green) and DsRed (red), with the DsRed images pseudo colored (ICA) to enhance the contrast of neuritic processes. Neuronal soma (white arrows) and axonal processes (black arrowheads) are indicated. Scale bars: 50 μm. **B**) Quantitative analysis of DsRed-positive axon length in YFP-ArfGAP1-positive neurons or control neurons (empty vector) is shown. Bars represent the mean ± SEM axon length (in μm) from 90-120 double DsRed-/YFP-ArfGAP1-positive neurons, or single DsRed-positive neurons (control), across at least three independent experiments/cultures. \*\*\**P*<0.001 or \*\*\*\**P*<0.0001 compared to control (DsRed alone), or as indicated, by one-way ANOVA with Dunnett’s multiple comparisons test. *ns*, non-significant.

### Blocking ArfGAP1 phosphorylation rescues G2019S LRRK2-induced neurite outgrowth inhibition

We have shown that the co-expression of ArfGAP1 and LRRK2 has a synergistic effect on reducing neurite length that is dependent in part on normal LRRK2 GTPase activity [16]. To evaluate the impact of ArfGAP1 phosphorylation sites on the interaction with LRRK2, we co-transfected cortical neurons with a combination of FLAG-tagged LRRK2 (WT or G2019S), YFP-ArfGAP1 variants (WT, 3A or 3D) and pDsRed-Max-N1 plasmids at a 10:10:1 molar ratio. Initially, we find that expression of G2019S LRRK2 or WT ArfGAP1 alone markedly and equivalently reduce neurite length by 35-40% compared to WT LRRK2 or control (empty vector) neurons (**Fig. S4**). The co-expression of WT or 3D ArfGAP1 with G2019S LRRK2 reduces neurite length by 50-60% compared to control neurons, whereas surprisingly the phospho-null 3A mutant dramatically protects from the effects of G2019S LRRK2 with only a 20-25% reduction of neurite length (**Fig. 5** and **S4**). Collectively, these data suggest that ArfGAP1 phosphorylation at these sites is required and sufficient for the synergistic interaction with G2019S LRRK2 in reducing neurite length, and blocking these sites (in the 3A mutant) can interfere with the neurite outgrowth deficits induced by G2019S LRRK2.

**Figure 5.**
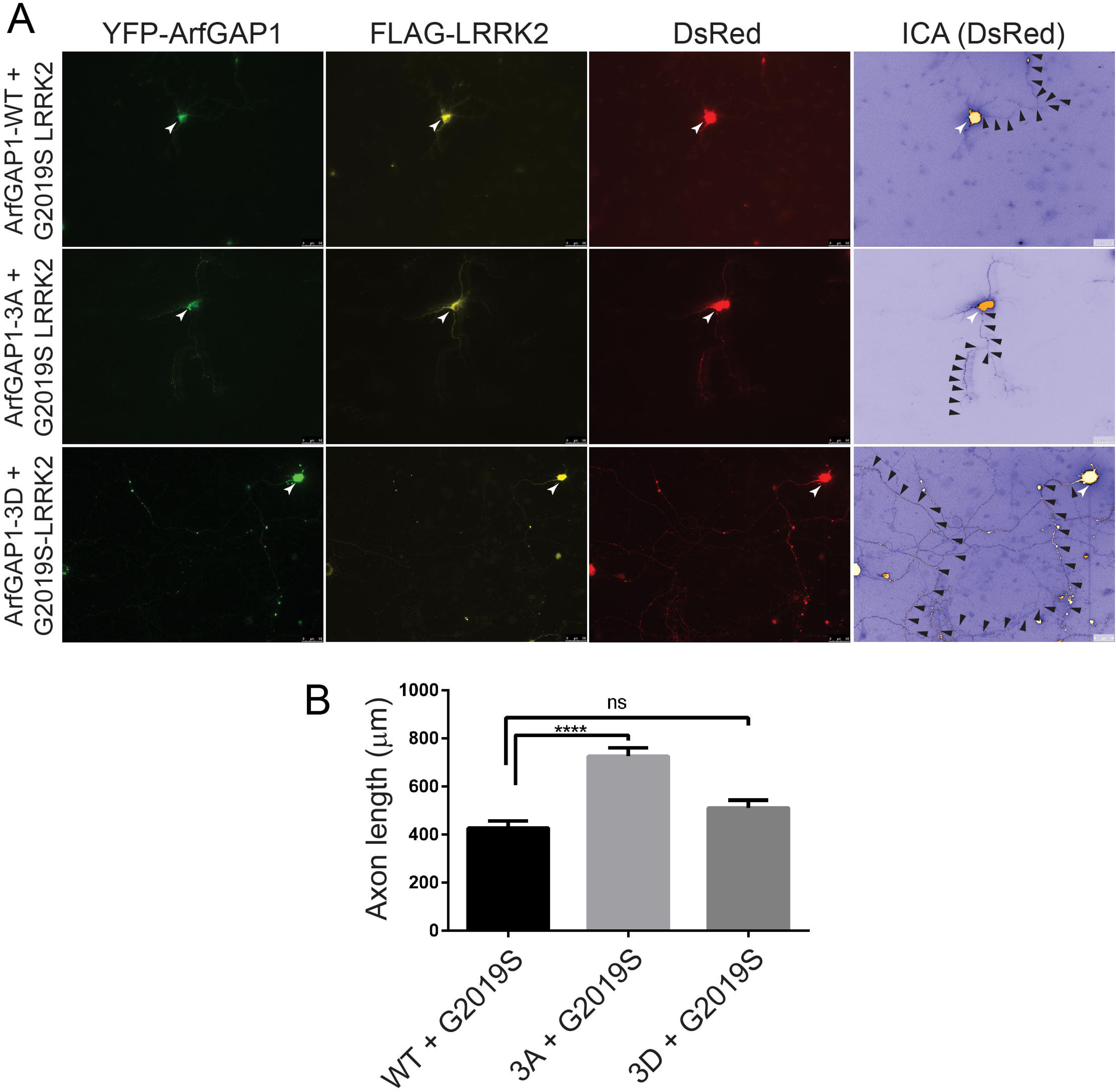
Preventing ArfGAP1 phosphorylation protects against the inhibition of neurite outgrowth induced by G2019S LRRK2. **A**) Rat primary cortical neurons were co-transfected at DIV3 with YFP-ArfGAP1 (WT, 3A or 3D), FLAG-LRRK2 (G2019S) and DsRed-Max-N1 plasmids, and fixed at DIV6 for immunofluorescence analysis with anti-FLAG antibody. Fluorescent images reveal the co-labeling of cortical neurons with YFP-ArfGAP1 (green), G2019S LRRK2 (yellow) and DsRed (red), with the DsRed images pseudo colored (ICA) to enhance the contrast of neuritic processes. Neuronal soma (white arrows) and axonal processes (black arrowheads) are indicated. Scale bars: 50 μm. **B**) Quantitative analysis of DsRed-positive axon length in YFP-ArfGAP1-/LRRK2-positive neurons is shown. Bars represent the mean ± SEM axon length (in μm) from 90-120 triple DsRed-/YFP-ArfGAP1-/LRRK2-positive neurons across at least three independent experiments/cultures. \*\*\*\**P*<0.0001 compared to WT ArfGAP1/LRRK2 by one-way ANOVA with Dunnett’s multiple comparisons test. *ns*, non-significant.

### ArfGAP1 interactome analysis identifies proteins localized to the Golgi, mitochondrial and cytoplasmic compartments

Since LRRK2-specific ArfGAP1 phospho-sites are localized within the non-catalytic domain, and specifically the ALPS2 motif, and regulate ArfGAP1-induced Golgi dispersal and neurite outgrowth deficits, we next sought to explore the protein interactome of ArfGAP1 and the potential impact of phosphorylation. Large-scale protein-protein interaction studies serve an important role in understanding the biological processes and cellular functions of a protein that are largely dependent on its subcellular localization and activity. ArfGAP1 can cycle between the Golgi complex, Golgi-derived vesicles and the cytosol [39], with phosphorylation impacting its actions at the Golgi. Accordingly, we first explored the interaction partners of ArfGAP1 in human cells by mass spectrometry. Detergent-soluble extracts derived from SH-SY5Y neural cells transiently expressing YFP-tagged wild-type ArfGAP1 were subjected to co-immunoprecipitation (IP) assays using anti-GFP antibody to enrich for interacting partners of ArfGAP1 and submitted for LC-MS/MS analysis. Similar anti-GFP IPs from cell extracts transfected with an empty plasmid served as a control to identify non-specific binding proteins that were subsequently excluded. MS-based proteome analysis with label-free protein quantification of biological triplicate experiments identifies 114 putative interacting proteins (FDR <0.05, fold change ≥ 4) including several known ArfGAP1-interacting partners such as adaptor protein complex AP-2 subunit alpha-1 (AP2A1) [47] and COPI coatomer subunits (COPA, COPB1, COPB2, COPG1) [48] (**Fig. 6A, Table S1**). The interactome screen also identifies several novel ArfGAP1 interactors, such as the mitochondrial proteins VDAC1, VDAC2, VDAC3 and TOMM40, as well as Rab1B, Rab5C and Rab11A that are localized to intracellular vesicular membranes (**Fig. 6A, Table S1**). Surprisingly, we do not identify Arf1 or LRRK2 in our interactome dataset, potentially due to their labile interaction and/or low abundance in SH-SY5Y cells.

**Figure 6.**
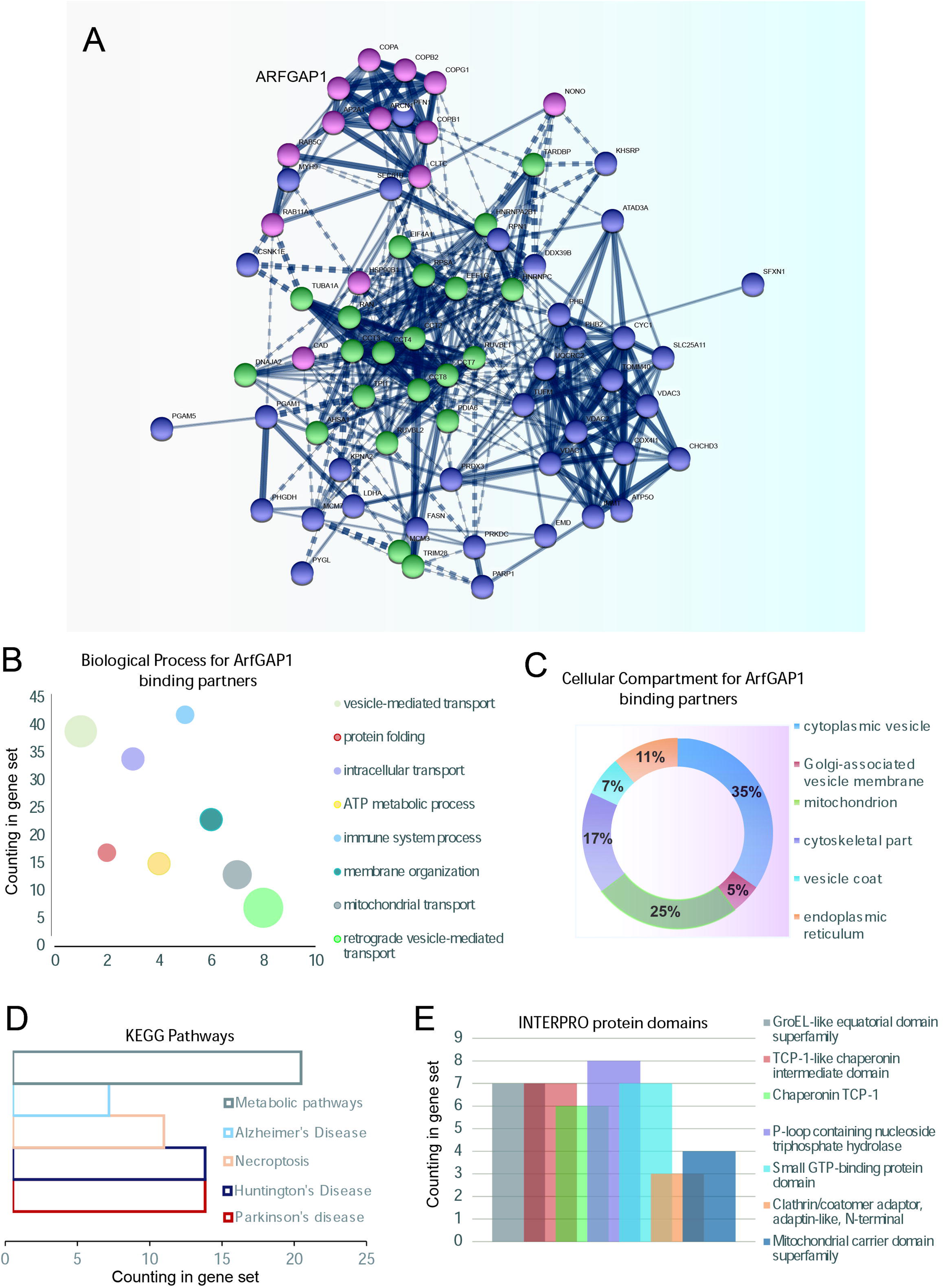
Protein interactome of ArfGAP1 reveals binding to a collection of mitochondrial proteins in neural cells. SH-SY5Y cells transiently expressing wild-type YFP-tagged ArfGAP1, or empty plasmid as a control, were subjected to immunoprecipitation with an anti-GFP antibody and subjected to LC-MS/MS analysis. **A**) K-means clustering of ArfGAP1 interaction maps of binding proteins identified by MS. Candidate interacting proteins (114) are segregated into three distinct clusters based on subcellular location: mitochondrial membrane (purple), cytoplasmic part (green) and Golgi membrane vesicle (pink). **B-C**) Gene Ontology (GO) analysis of ArfGAP1-interacting partners. **B**) GO biological process indicates ∼40 interacting proteins involved in “vesicle-mediated transport”, whereas **C**) Pie chart represents the GO cellular compartments of all binding partners. **D**) KEGG pathways associated with ArfGAP1 interactors. **E**) INTERPRO protein domain analysis of ArfGAP1 interactors.

To explore the intermolecular connections between all identified binding partners, we integrated Search Tool for Retrieval of Interacting Genes (STRING) analysis with our MS-based ArfGAP1 interactome data. The KMEANS clustering in protein-protein interaction networks identifies three distinct clusters, i) mitochondrial membrane, ii) cytoplasmic part and iii) Golgi membrane vesicle, suggesting that ArfGAP1-interacting partners generally segregate into these three subcellular compartments (**Fig. 6A**). To derive additional insight from our interactome data, Gene Ontology (GO) analysis was conducted to describe the biological processes and cellular compartments of ArfGAP1-interacting partners. For GO biological processes, 39 proteins are linked to vesicle-mediated transport, whereas there are also 6 proteins (COPA, COPB1, COPB2, COPG1, ARCN1 and Rab1B) specifically linked to retrograde vesicle-mediated transport (**Fig. 6B**). For GO cellular compartments, most interactors (35%) are cytoplasmic vesicle proteins with only 5% of proteins localizing to Golgi-associated vesicle membranes, whereas surprisingly 25% of proteins are associated with mitochondria (**Fig. 6C**). We also performed pathway analysis on the ArfGAP1 interactome using the KEGG Pathways database. Notably, this analysis identifies “PD pathways” along with four other pathways including “Huntington’s disease” and “Alzheimer’s disease” (*P*<0.05) (**Fig. 6D**), and there are several interacting partners, such as VDAC1-3 that are directly linked with PD pathways [49]. We further attempted to classify interacting proteins into protein families to potentially identify key domains within the ArfGAP1-interacting protein network. INTERPRO analysis predicts several protein domains, including the mitochondrial carrier domain superfamily, P-loop containing nucleoside triphosphate hydrolase, and small GTP-binding protein domains, that are shared by at least 20 interacting partners. Of note, five interactors contain a small GTP-binding protein domain (Rab1B, Rab5C, Rab11A, Ran and EEF1G). Taken together, our comprehensive MS-based interactome data provides a global view of ArfGAP1 interaction networks and also provides an unexpected connection to mitochondrial membranes.

### ArfGAP1 phosphorylation impacts its subcellular localization and interaction with VDACs

From the ArfGAP1 interactome analysis, the connection to mitochondrial membranes was both intriguing and unappreciated in prior studies. As such, voltage-dependent anion channels (VDAC) 1-3, that are localized to the outer mitochondrial membrane, emerged as interesting and novel interacting partners of ArfGAP1. We elected to evaluate whether ArfGAP1 phosphorylation influences the interaction with VDACs by conducting co-IP assays with anti-GFP antibody from SH-SY5Y cell extracts transiently expressing YFP-tagged ArfGAP1 variants (WT, 3A or 3D). Intriguingly, the phospho-mimic 3D mutant markedly increases the interaction between ArfGAP1 and VDACs (using a pan-VDAC1-3 antibody) compared to WT or 3A ArfGAP1 (**Fig. 7A**). As VDACs are integral transmembrane proteins that traverse the outer mitochondrial membrane [50], we hypothesized that the ArfGAP1-3D variant is in close proximity to mitochondria. We therefore conducted confocal fluorescence microscopy on SH-SY5Y cells transiently expressing YFP-ArfGAP1 variants together with the outer mitochondrial membrane marker, TOMM20. Correlation analysis reveals a significant increase in the colocalization of ArfGAP1-3D with mitochondrial TOMM20, compared to WT or 3A ArfGAP1 (**Fig. 7B**). While ArfGAP1 is predominantly a Golgi-localized protein [51–53], ArfGAP1-3D is mostly cytoplasmic whereas WT and 3A ArfGAP1 adopt a more typical perinuclear localization (**Fig. 7B**) [51–53]. As the C-terminal domain is involved in regulating the subcellular localization of ArfGAP1 and its interaction with membranes [51, 53], particularly via its ALPS 1 and 2 motifs, we asked whether the phospho-null and phospho-mimic variants adopt different localizations. Confocal fluorescence analysis of SH-SY5Y cells expressing YFP-ArfGAP1 variants reveals that WT ArfGAP1, and to a greater extent the 3A mutant, mostly co-localize with the GM130-labeled Golgi complex, whereas ArfGAP1-3D adopts a distinct cytoplasmic-like localization (**Fig. 8A**). To confirm this differential localization of the phospho-mimic 3D mutant, we performed subcellular fractionation analysis on SH-SY5Y cells expressing YFP-ArfGAP1 variants. We detect WT ArfGAP1 broadly distributed across all subcellular fractions, with localization to nuclear, microsomal (i.e., Golgi), cytoplasmic and mitochondrial fractions (**Fig. 8B**). We find that ArfGAP1-3A is significantly enriched in the microsomal fraction, whereas ArfGAP1-3D is significantly enriched in the cytoplasmic fraction, compared to WT ArfGAP1 (**Fig. 8B**). ArfGAP1 variants display similar levels of enrichment in the nuclear and mitochondrial fractions (**Fig. 8B**). These fractionation data confirm the differential localization of 3A and 3D ArfGAP1 mutants indicated by immunofluorescence analysis (**Fig. 8A**) and suggest that the phospho-null 3A mutant drives a Golgi localization whereas the phospho-mimic 3D mutant drives a cytoplasmic localization. Collectively, our data suggests that phosphorylation at these sites regulates ArfGAP1 subcellular localization between the Golgi and cytoplasmic compartments, and the interaction with outer mitochondrial membrane proteins such as VDACs.

**Figure 7.**
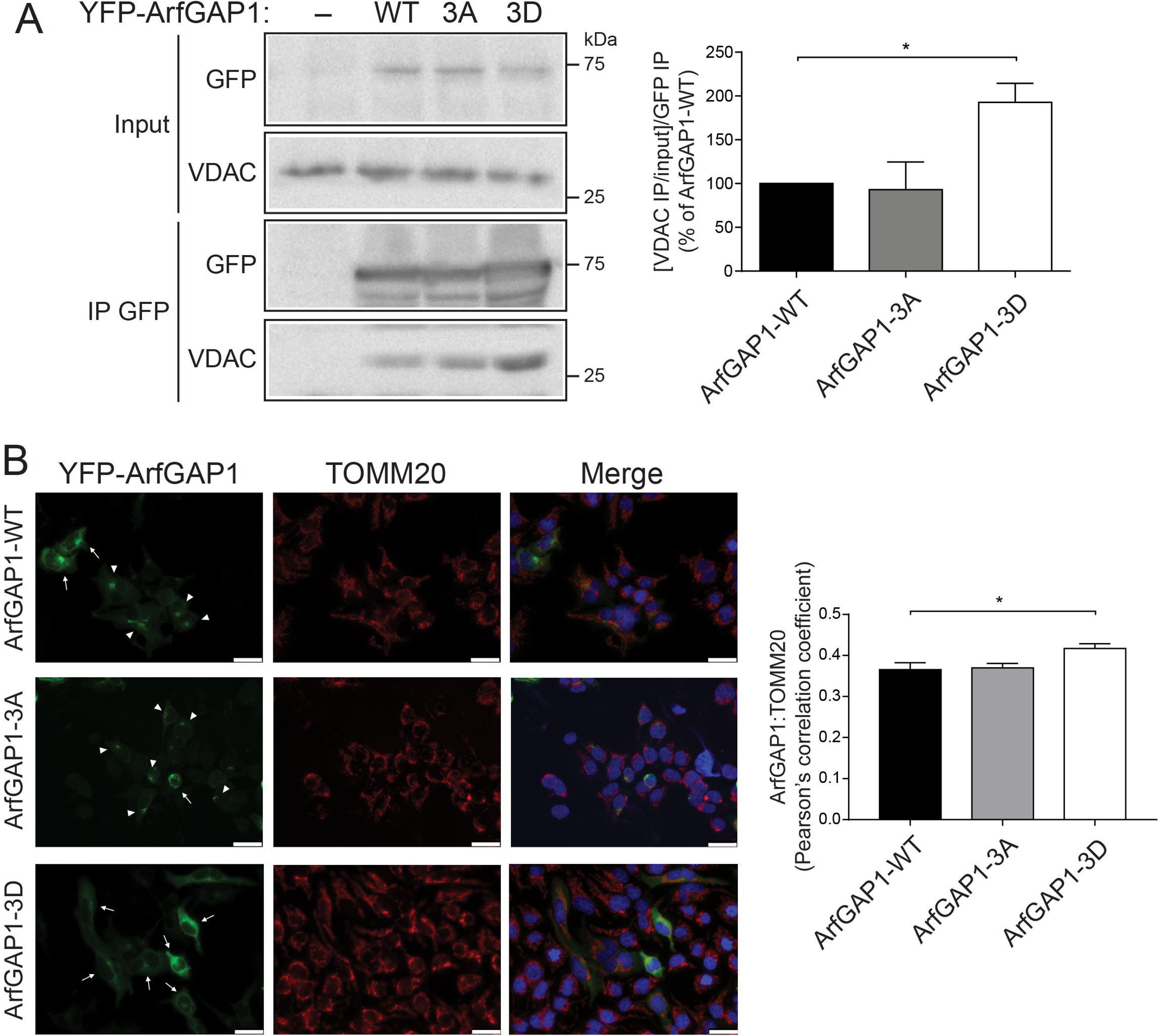
Phospho-mimic mutations increase ArfGAP1 binding to VDACs and its mitochondrial localization. **A**) SH-SY5Y cell extracts transiently expressing YFP-ArfGAP1 (WT, 3A or 3D) or empty plasmid were subjected to immunoprecipitation (IP) with anti-GFP antibody, and IP and input lysates were probed by Western blotting with anti-pan-VDAC1-3 and anti-GFP antibodies. Graph indicates densitometric analysis of VDAC binding for each ArfGAP1 variant, with data expressed as IP VDAC normalized to input VDAC levels, and ratios further normalized to IP ArfGAP1 levels. Bars represent mean ± SEM (*n* ≥ 3 independent experiments). Molecular mass is indicated in kilodaltons (kDa). **B**) SH-SY5Y cells expressing YFP-ArfGAP1 (WT, 3A or 3D) subjected to immunofluorescence with anti-TOMM20 antibody. Representative fluorescent microscopic images indicate partial colocalization of mitochondrial TOMM20 (red) and YFP-ArfGAP1 (green). Nuclear DAPI (blue) is shown in merged images. Arrows indicate diffuse localization of YFP-ArfGAP1 (mainly 3D mutant) and arrowheads indicate perinuclear Golgi localization (mainly WT and 3A mutant). Scale bars: 25 µm. Graph indicates Pearson’s correlation coefficients for ArfGAP1/TOMM20 colocalization. Bars represent mean ± SEM (*n* ≥ 3 independent experiments). \**P*<0.05 compared to WT ArfGAP1 by one-way ANOVA with Dunnett’s multiple comparisons test, as indicated.

**Figure 8.**
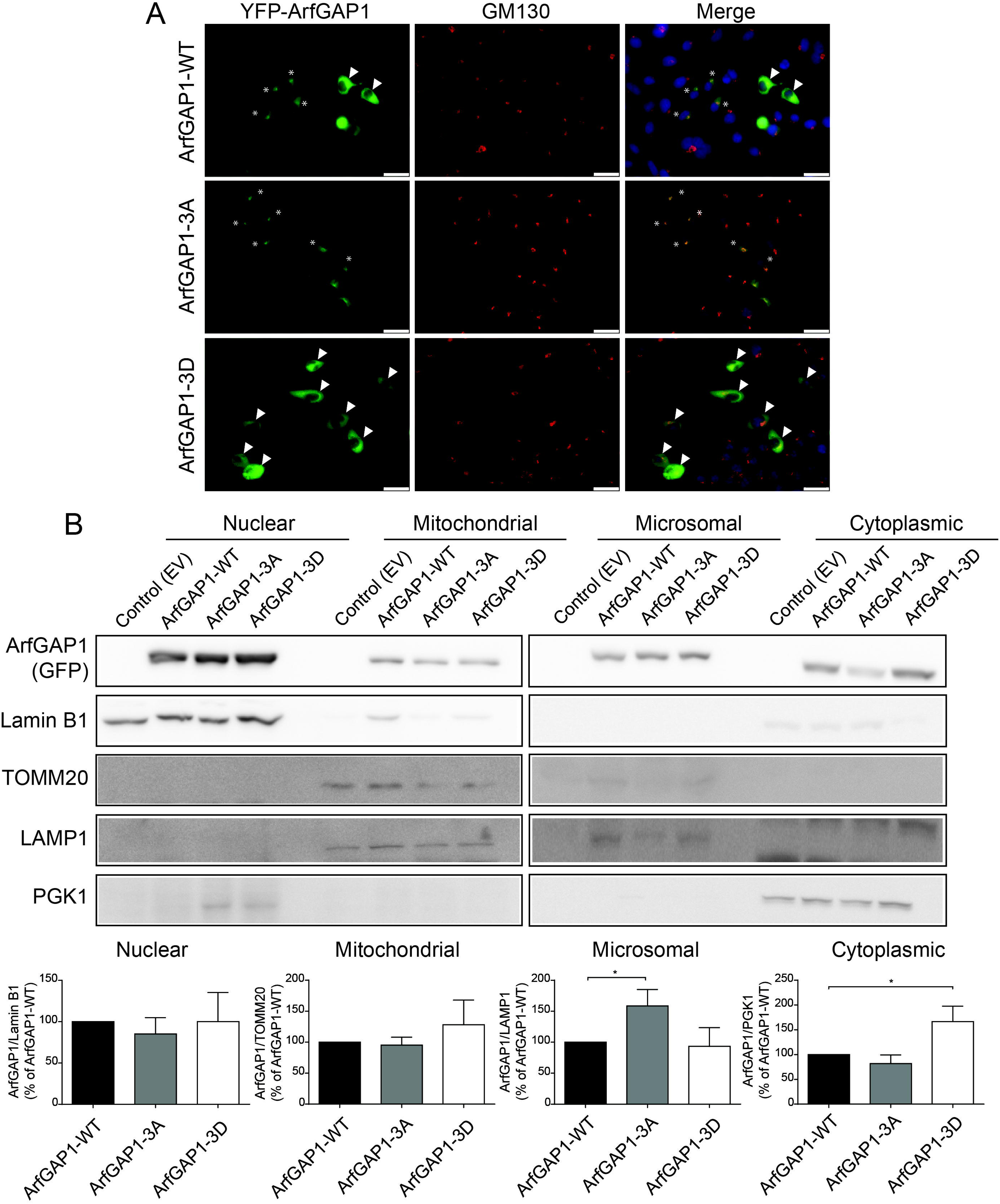
Phospho-mimic mutations disrupt the localization of ArfGAP1 to the *cis*-Golgi and drive a cytoplasmic location. **A**) SH-SY5Y cells expressing YFP-ArfGAP1 (WT, 3A or 3D) subjected to immunofluorescence with anti-GM130 antibody. Representative fluorescent microscopic images indicate colocalization of *cis*-Golgi GM130 (red) and YFP-ArfGAP1 (green). Nuclear DAPI (blue) is shown in merged images. Asterisks indicate perinuclear Golgi localization of YFP-ArfGAP1 (3A mutant > WT) and arrowheads indicate diffuse cytoplasmic localization (3D mutant >> WT). Scale bar: 25 µm. **B**) Subcellular fractionation by differential centrifugation of SH-SY5Y cell extracts expressing YFP-ArfGAP1 (WT, 3A or 3D) or control empty vector. Western blot analysis of nuclear, mitochondrial, microsomal and cytoplasmic fractions with anti-GFP antibody and antibodies to markers enriched in each fraction (Lamin B1/nuclear, TOMM20/mitochondria, LAMP1/microsomes, PGK1/cytoplasmic). ArfGAP1 variants are broadly detected across the different subcellular fractions, with subtle differences. Graphs indicate densitometric analysis of YFP-ArfGAP1 levels in each subcellular fractionation normalized to their respective compartment marker. Bars represented the mean ± SEM levels of ArfGAP1 (*n* ≥ 3 independent experiments). \**P*<0.05 compared to WT ArfGAP1 by one-way ANOVA with Dunnett’s multiple comparisons test, as indicated.

### ArfGAP1 phosphorylation regulates the formation of Golgi-derived vesicles induced by ER stress

As ArfGAP1 is involved in Golgi-to-ER retrograde transport [38], we asked whether the altered localization of ArfGAP1 due to phosphorylation can influence the cellular response and Golgi sorting induced by ER stress. As such, SH-SY5Y cells transiently co-expressing YFP-ArfGAP1 variants (WT, 3A or 3D) and mCherry-N1-galactosyltransferase (GalT), a Golgi-resident enzyme, were treated with or without tunicamycin (2.5 µg/ml) for 3 h. Under basal conditions (DMSO treatment), the small number of cells displaying GalT-positive Golgi-derived vesicles are similar between control (empty vector) or WT ArfGAP1 conditions (**Fig. 9**). ArfGAP1-3A expression markedly increases the number of cells with GalT-positive vesicles localized outside the Golgi complex (∼20%), whereas ArfGAP1-3D has an intermediate, yet non-significant effect compared to WT ArfGAP1 (**Fig. 9**). Interestingly, while tunicamycin treatment that induces mild ER stress has no effect on the number of cells with GalT-positive vesicles in the absence of ArfGAP1, the expression of WT or 3A ArfGAP1 produces a marked and equivalent increase in cells displaying GalT-positive vesicles (40-50%) (**Fig. 9**). In contrast, ArfGAP1-3D has a small effect on the number of cells with GalT-positive vesicles following tunicamycin treatment, similar to its effects in DMSO-treated cells (**Fig. 9**). Our data indicate that ArfGAP1 expression robustly induces the formation of GalT-positive Golgi-derived vesicles in response to mild ER stress, consistent with an increase in Golgi-to-ER retrograde transport, whereas the phospho-mimic 3D mutant completely inhibits the normal activity of ArfGAP1 in this assay. These data are consistent with the altered cytoplasmic localization of ArfGAP1-3D compared to the WT and 3A proteins (**Fig. 8**).

**Figure 9.**
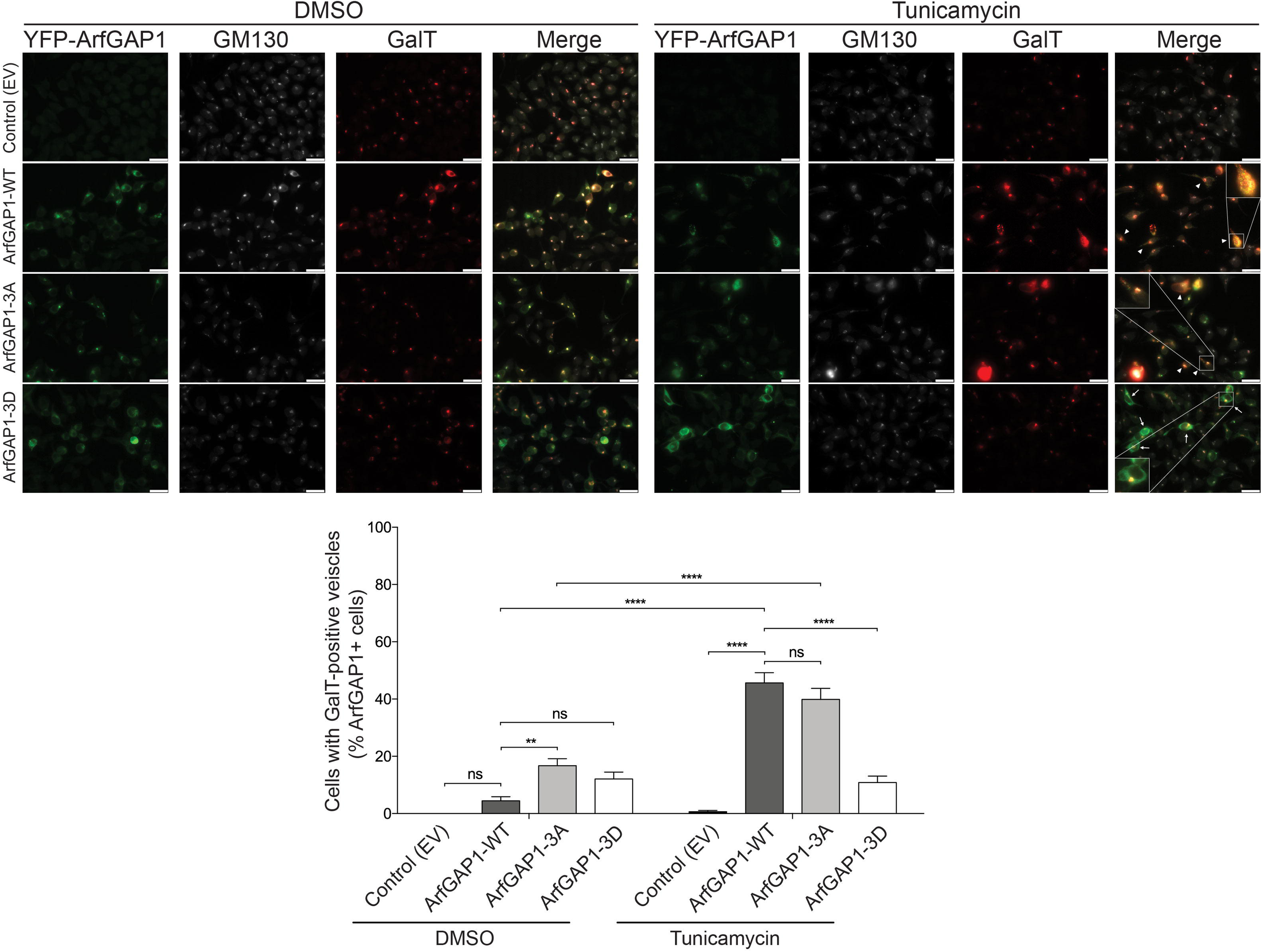
Phospho-mimic mutations disrupt ArfGAP1-induced Golgi-derived vesicle formation under mild ER stress. SH-SY5Y cells transiently co-expressing YFP-ArfGAP1 (WT, 3A, 3D) and mCherry-GalT were treated with vehicle (DMSO) or tunicamycin (2.5 µg/ml) for 3 h and subjected to immunofluorescence with an anti-GM130 antibody. Representative fluorescent microscopic images indicate colocalization of GalT (red) and *cis*-Golgi GM130 (white) in YFP-ArfGAP1-positive cells (green), whereas GalT signal often extends beyond the perinuclear GM130-positive Golgi complex. Presence (arrowhead) or absence (arrow) of GalT-positive Golgi-derived vesicles are observed in YFP-ArfGAP1 overexpressing cells. Scale bar: 25 µm. Graph indicates percent of ArfGAP1-positive cells with GalT-positive vesicles (dispersed outside of the Golgi complex) following tunicamycin (or DMSO) treatment. Bars represent the mean ± SEM percent of cells with GalT-positive vesicles (*n* ≥ 3 independent experiments). \*\**P*<0.01 or \*\*\*\**P*<0.0001 by two-way ANOVA with Tukey’s post-hoc analysis, as indicated. *ns*, non-significant.

## Discussion

In this study we have evaluated the LRRK2-ArfGAP1 interaction, with a focus on the impact of LRRK2-specific phosphorylation on ArfGAP1 localization and activity. LRRK2 robustly and directly phosphorylates ArfGAP1 *in vitro*, with the most prominent phosphorylation sites residing within C-terminal residues 252-359 of ArfGAP1. Mass spectrometry analysis identified four phosphorylated threonine residues (pThr145, pThr189, pThr230, pThr292) in full-length ArfGAP1, with only one of these sites (pT292) localized within residues 252-359. A prior study identified additional LRRK2-dependent phosphorylation sites, including Ser284 within this region [17], that were also incorporated into our study together with a known phosphorylation site at T291. Combining null mutations at these three sites (S284A, T291A, T292A) is sufficient to block LRRK2-mediated phosphorylation within residues 252-359 of ArfGAP1. Using phospho-null (Ala, A) or phospho-mimic (Asp, D) substitutions combined at these three sites, we find that 3A and 3D mutations both impair ArfGAP1-induced Golgi complex fragmentation in human neural cells. We further find that blocking phosphorylation at these sites (3A mutant) is sufficient to attenuate neurite outgrowth deficits normally induced by ArfGAP1 in cultured primary neurons and rescues the pathogenic effects of G2019S LRRK2 on neurite outgrowth. We identify novel interacting proteins for ArfGAP1 in human neural cells, including mitochondrial proteins, and demonstrate that the phospho-mimic 3D mutation promotes the interaction of ArfGAP1 with mitochondrial VDACs. The enhanced interaction with VDACs most likely results from the increased cytoplasmic localization of the ArfGAP1-3D mutant and its impaired localization to the Golgi complex. Consistent with this altered localization, the 3D mutant completely blocks the ArfGAP1-induced formation of Golgi-derived vesicles following mild ER stress. Collectively, our study reveals the functional impact of selected LRRK2-specific phosphorylation sites within ArfGAP1 and identifies the opposing actions of ArfGAP1 phosphorylation in i) maintaining Golgi localization and promoting Golgi complex dispersal (impaired by the phospho-mimic mutant) and in ii) mediating neurite outgrowth deficits (rescued by the phospho-null mutant). Our findings indicate that ArfGAP1 subcellular localization, protein interactions and activity are dynamically regulated by LRRK2-specific phosphorylation, suggesting that the interaction between LRRK2 and ArfGAP1 is complex and non-linear with phosphorylation serving to regulate different aspects of ArfGAP1 function.

Our prior studies with ArfGAP1 demonstrated that its expression was critical for LRRK2-induced cellular toxicity in yeast or primary neuronal models [16, 31], implying that ArfGAP1 normally plays a role in mediating the downstream pathogenic effects of mutant LRRK2. However, it was unclear whether such an effect results from the GAP-like activity of ArfGAP1 that enhances the GTP hydrolysis activity of LRRK2 (and other small GTPases), or whether it results from the direct phosphorylation of ArfGAP1 by LRRK2 and its downstream consequences on Golgi vesicle trafficking pathways [16, 17]. As a potential mechanism for explaining LRRK2-induced neurotoxicity, we have explored the role of ArfGAP1 as a LRRK2 substrate. A prior study suggested that mutation of six distinct serine or threonine residues located throughout ArfGAP1 completely prevented its phosphorylation by LRRK2, but the relative contribution of phosphorylation at each site was not clearly determined [17]. Here, by combining domain mapping and mutational studies, we have highlighted residues 252-359 as a major region for ArfGAP1 phosphorylation by LRRK2 and nominated three candidate phosphorylation sites in this region (S284, T291 and T292). These phosphorylation sites are located within a highly conserved lipid-packing sensor motif (ALPS2, residues 264-295) that is known to regulate the binding of ArfGAP1 to curved membranes [44, 45]. Notably, we also identified another phosphorylation site (pT230) within the ALPS1 motif (residues 199-234), not further studied here, suggesting a more general role for LRRK2 in regulating ArfGAP1 membrane association. Consistent with such a function, phospho-mimicking mutations at the ALPS2 sites (S284D/T291D/T292D) induced the relocalization of ArfGAP1 from the *cis*-Golgi compartment to the cytoplasm, where it exhibits an enhanced interaction with VDAC proteins at the mitochondrial outer membrane. The 3D mutation also impaired ArfGAP1-induced Golgi fragmentation and impaired the ER stress-induced production of Golgi-derived vesicles, all consistent with a diminished association of ArfGAP1-3D with the *cis*-Golgi membrane. Prior studies have shown that phosphorylation within the C-terminal domain re-localizes ArfGAP1 from the Golgi to the cytosol [53]. Therefore, we suggest that LRRK2-mediated phosphorylation within the ALPS2 motif alters its affinity or specificity for curved membranes of organelles. A parallel role has been ascribed to the LRRK2-mediated phosphorylation of endophilin-A in endocytosis, dynamically affecting its membrane association and tubulation via Ser75 phosphorylation within its membrane-sensing BAR domain [54]. The ArfGAP1-3A mutant also impairs ArfGAP1-induced Golgi fragmentation yet adopts a normal subcellular localization at the *cis*-Golgi compartment. It is unclear whether these effects result from altered membrane affinity or dynamics, or perhaps from altered GAP activity directed at Golgi-localized small GTPases such as Arf1. While we have not directly tested the impact of ALPS2 motif phosphorylation on GAP activity, especially since the GAP domain is distant and oppositely resides in the N-terminal region of ArfGAP1, a prior study demonstrated that LRRK2-mediated phosphorylation impaired ArfGAP1-stimulated Arf1 GTP hydrolysis *in vitro* [17]. However, a non-phosphorylatable 6A mutant of ArfGAP1 retained the capacity to stimulate Arf1 GTPase activity *in vitro*, implying that it was fully functional [17]. Accordingly, in cells, we propose that ArfGAP1-3A does not impact its GAP activity but instead exerts its effects by altered membrane binding.

ALPS motifs play a central role in sensing membrane curvature, and target proteins onto these membranes depending on the degree of curvature [44, 53]. ALPS are amphipathic helices where the hydrophobic-face residues adsorb to membranes that display lipid packing defects, and with a polar face enriched in serine and threonine residues [44, 45]. ArfGAP1 contains two ALPS motifs [45]. Interestingly, these ALPS motifs present differences regarding membrane curvature and lipid composition to which they bind, with ALPS1 binding to highly curved membranes whereas ALPS2 appears to tune this detection toward larger radii curved-membranes [45, 55]. A previous study demonstrated that substitution of amino acids at the hydrophobic face of ALPS1 can modify ArfGAP1 cellular localization [51]. Here, we present evidence suggesting that the ArfGAP1-VDAC interaction could be regulated by LRRK2-dependent phosphorylation. This increased interaction does not imply that the ALPS2 motif interacts with VDAC, but rather we suggest that the shift between high and low curvature membrane detection induced by ALPS2 phosphorylation localizes ArfGAP1 to the outer mitochondrial membrane, thereby facilitating an ArfGAP1-VDAC interaction. Pathogenic LRRK2 mutants are associated with mitochondrial dysfunction (extensively reviewed in [56]), although it is debated whether LRRK2 affects mitochondria directly or indirectly. Previous studies have shown that cytosolic proteins can regulate VDAC function and affect mitochondrial outer membrane permeability [57, 58]. Our data demonstrates that ArfGAP1 phosphorylation (induced by LRRK2) increases its interactions with VDAC and allows us to speculate that ArfGAP1 phosphorylation may play a pathogenic role in PD by regulating mitochondrial function. Additional experiments exploring the ArfGAP1-VDAC interaction will be required to support such a hypothesis.

Since ALPS motifs bind to membranes in a specific manner recognizing lipid packaging defects, it is interesting to consider ALPS phosphorylation as a functional regulatory mechanism for ArfGAP1 localization. Moreover, as the lipid composition of membranes varies between different organelles, it is plausible that ArfGAP1 phosphorylation by LRRK2 will modify its affinity for membranes with different curvature and/or lipid composition, thereby regulating ArfGAP1 localization and its role within different intracellular processes [59]. Such a mechanism for protein regulation has already been observed in yeast, where Vps41 can alternate between different trafficking routes depending on phosphorylation of its ALPS motif, suggesting phosphorylation could serve as a switch for Vps41 between endosome-to-vacuole and AP-3 vesicle-to-vacuole fusion [60–62]. Adaptor proteins (AP) are key players in vesicular trafficking process between cellular compartments, connecting endosomes, Golgi, plasma membrane and lysosomes [63, 64]. The interaction of ArfGAP1 with AP-2 and AP-3 has already been demonstrated, suggesting that ArfGAP1 is likely involved in other intracellular pathways in addition to retrograde Golgi-ER transport [64–66]. Accordingly, Meng and colleagues recently showed that ArfGAP1 binds to mTORC1, inhibiting its activation and promoting autophagy activation [67]. Together, ALPS phosphorylation is likely to influence ArfGAP1 localization and intracellular functions by regulating which proteins and membranes it can interact with.

The capacity of ArfGAP1 expression to modulate neurite outgrowth in cultured primary neurons suggests a potential role in regulating neuronal process complexity or integrity. However, there are limited studies on ArfGAP1 in the context of brain localization, structure or function, especially due to a paucity of specific antibodies [51, 68]. Our prior study demonstrated that endogenous ArfGAP1 is detected in multiple brain regions and is localized within different primary neurons, and WT ArfGAP1 overexpression was sufficient to inhibit neurite outgrowth, similar to PD-linked LRRK2 mutants [16]. Here, we show that blocking LRRK2 phosphorylation via the 3A mutant impairs the capacity of ArfGAP1 to regulate neurite outgrowth. This might imply that ArfGAP1 phosphorylation *per se* is required for this activity, as suggested by the similar effects of WT and phospho-mimic 3D ArfGAP1. Consistent with this idea, we have previously shown that endogenous LRRK2 is required, at least in part, for the effects of ArfGAP1 on neurite outgrowth [16]. Our data also reveals that 3A ArfGAP1 is able to rescue the robust effects of G2019S LRRK2 on neurite outgrowth inhibition. The mechanism underlying this interesting rescue effect is unclear but it could be due to interference with the normal phosphorylation of endogenous ArfGAP1 perhaps via sequestration of, or excessive binding to, LRRK2. Supporting this idea, we have previously shown that reducing ArfGAP1 expression is sufficient to protect against G2019S LRRK2 in these neurite assays [16]. It is therefore tempting to speculate that non-phosphorlyatable ArfGAP1 peptides could represent an interesting strategy for attenuating the neurotoxic effects of mutant LRRK2 in future studies.

Collectively, the present study together with prior work [16, 17, 31] provides further evidence for a functional interaction between LRRK2 and ArfGAP1 that serves to regulate ArfGAP1 subcellular localization, protein interactions, activity and neuronal toxicity via LRRK2-mediated phosphorylation of its ALPS2 motif. Our data suggests a complex relationship between ArfGAP1 and LRRK2 in the pathogenesis of PD, linking Golgi-ER retrograde sorting and mitochondrial pathways through ArfGAP1 phosphorylation. Our findings support additional validation of ArfGAP1 as a putative therapeutic target for modulating *LRRK2*-linked PD.

## Materials and methods

### Animals

Timed pregnant female Sprague-Dawley outbred rats were obtained from Taconic Biosciences and P1 rats were used to prepare post-natal primary cortical neuronal cultures as previously reported [16]. Mice were maintained in a pathogen-free barrier facility and provided with food and water *ad libitum* and exposed to a 12 h light/dark cycle. Animals were treated in strict accordance with the NIH Guidelines for the Care and Use of Laboratory Animals. All animal experiments were approved by the Van Andel Institute Institutional Animal Care and Use Committee (IACUC).

### Expression plasmids, proteins and antibodies

Mammalian expression plasmids containing FLAG-tagged full length human LRRK2 (WT, R1441C and G2019S) variants were kindly provided by Dr. Christopher Ross (Johns Hopkins University, Baltimore, USA) [69]. A plasmid encoding mCherry-N1-Galactosyltransferase (GalT) was obtained from Addgene (#87327) [70]. A plasmid containing DsRed-Max-N1 was obtained from Addgene (#21718) [71]. A C-terminal YFP-tagged rat ArfGAP1 plasmid was kindly provided by Dr. Jennifer Lippincott-Schwartz (National Institutes of Health, Bethesda, USA) [39]. ArfGAP1 phospho-null/phospho-mimic mutations (S284A, S284D, T291A, T291D, T292A, T292D, S284A/T291A/T292A and S284D/T291D/T292D) were generated by site-directed mutagenesis using the Stratagene QuickChange II XL kit (Agilent Technologies, La Jolla, CA, USA) and DNA sequencing was performed to confirm integrity.

Recombinant GST-tagged human LRRK2 proteins (residues 970-2527) were obtained from Invitrogen (Carlsbad, CA, USA). GST-tagged rat ArfGAP1 plasmids (full-length or deletion mutants [residues 1-415, 1-136, 137-251, 252-359, 137-415 and 360-415] in pGEX-6P1) were kindly provided by Dr. Victor Hsu (Brigham and Women’s Hospital, Harvard Medical School) [65]. Phospho-null variants (S284A, T291A, T292A and S284A/T291A/T292A) were introduced into a 252-359 (F5) GST-ArfGAP1 plasmid by site-directed mutagenesis. GST-ArfGAP1 proteins were purified from IPTG-induced bacteria using Glutathione-Sepharose 4B columns (GST Bulk Kit, Cytiva) following standard protocols as described [72].

The following primary antibodies were employed: mouse anti-FLAG (clone M2; Sigma-Aldrich, Buchs, Switzerland), mouse anti-GFP (clones 7.1 and 13.1; Roche Applied Science, Basel, Switzerland), mouse anti-GM130 (clone 35; BD Biosciences), goat anti-GST-HRP (RPN1236, GE Healthcare), mouse anti-TOMM20 (ab56783, Abcam), mouse anti-PGK1 (459250, Invitrogen), rabbit anti-LAMP1 (ab24170, Abcam), rabbit anti-Lamin B1 (ab16048, Abcam), and rabbit anti-pan-VDAC1-3 (PA1-954A, Thermo Fisher). Secondary antibodies included: HRP-coupled anti-mouse and anti-rabbit IgG, light chain-specific (Jackson ImmunoResearch, Inc., West Grove, PA, USA), and anti-rabbit IgG and anti-mouse IgG coupled to AlexaFluor-488, -546 and -633 (Invitrogen).

### Cell culture, transfection and treatment

Human SH-SY5Y neural cells were maintained at 37°C with 5% CO_2_ atmosphere in Dulbecco’s Modified Eagle’s Media (DMEM) (Gibco) supplemented with 10% (v/v) fetal bovine serum and penicillin/streptomycin. Cell transfection was performed using plasmid DNA and XtremeGene HP DNA Transfection reagent (Roche), according to manufacturer’s instructions. For endoplasmic reticulum stress assays, 48 h after co-transfection, cells were treated with tunicamycin (2.5 µg/ml) or DMSO for 3 h, followed by immunocytochemical analysis. Primary cortical neurons were maintained in 35 mm dishes on glass coverslips in Neurobasal media containing B27 supplement (2% w/v), L-glutamine (500 mM) and penicillin/streptomycin (100 U/ml) as previously described [16].

### Immunocytochemistry

SH-SY5Y neural cells were transiently transfected with YFP-ArfGAP1 variants, or co-transfected with YFP-ArfGAP1 and mCherry-N1-Galactosyltransferase (GalT) (Addgene #87327) at a 3:1 molar ratio. After 48 h, medium was removed, and cells were fixed with 4% PFA at room temperature for 20 min and then washed 3 times with 1X PBS (pH 7.4). Non-specific binding was blocked by incubation in 5% BSA in PBS at room temperature. Cells were incubated overnight at 4°C with anti-TOMM20 (Abcam) or anti-GM130 (BD Biosciences) antibodies, followed by incubation with AlexaFluor-conjugated secondary antibodies at room temperature for 90 min. Coverslips were mounted onto glass slides using Prolong Diamond Antifade Mountant with DAPI (Thermo Fisher). Fluorescence images were acquired by microscopy using a Nikon A1plus-RSi scanning confocal microscope or a Leica DM5500B epifluorescence microscope.

### Subcellular fractionation

For subcellular fragmentation, we followed the protocol from Abas and Luschnig [73] with some modifications. Culture medium was removed, cells were carefully washed once with PBS and resuspended in extraction buffer (EB): 100 mM Tris-HCl pH 7.5, 25% (w/w) sucrose, 5% (v/v) glycerol, 10 mM EDTA, 5 mM KCl, supplemented with Complete Mini Protease inhibitor EDTA-free cocktail (Roche) and Phosphatase Inhibitor cocktails 2 and 3 (Sigma Aldrich). Cells were homogenized using a 25G needle at 4°C, cell lysis was monitored with trypan blue, and samples were centrifuged once lysis was >95%. Briefly, lysates were centrifuged for 5 min at 800*g*. The nuclear fraction (pellet 1) was washed 2 times using EB. Supernatant was centrifuged for 20 min at 10,000*g*, and the mitochondrial fraction (pellet 2) was washed 2 times with EB. Supernatant was centrifuged for 120 min at 21,100*g*, the microsomal fraction (pellet 3) was washed twice with EB, and supernatant was saved as the cytosolic fraction. Samples were subjected to Western blot analysis.

### Co-Immunoprecipitation

For co-immunoprecipitation (IP) assays, SH-SY5Y cells were transiently transfected with the desired amount of plasmid in 10 cm dishes. At 48 h post-transfection, media was removed, and cells were harvested in 1 ml of lysis buffer (1X PBS, 1% Triton X-100, 1X Complete Mini Protease inhibitor cocktail [Roche]). Cell lysates were allowed to rotate 2 h at 4°C, and then centrifuged for 15 min at 15,000 rpm at 4°C. Supernatants were incubated overnight at 4°C with 50 µl Protein G-Dynabeads (Thermo Fisher) that had been pre-incubated for 2 h with anti-GFP antibody (2 µg; Roche). Samples were washed 3 times with 1X PBS, 1% Triton X-100 and once with PBS. IPs were eluted in 2X Laemmli buffer at 95°C for 5 min and analyzed by Western blot.

### Western blot analysis

Samples from IPs, input lysates, or subcellular fractionations were subjected to SDS-PAGE and transferred to nitrocellulose membranes (0.2 µm; GE Healthcare). Membranes were blocked for 1 h with 5% (w/v) nonfat milk and 0.1% Tween 20 in Tris-Buffered Saline (TBS-T) and incubated overnight at 4°C with primary antibodies: anti-FLAG, anti-GST-HRP, Anti-GM130, anti-VDAC, anti-GFP, anti-TOMM20, anti-PGK1, anti-LAMP1 or anti-Lamin B1. After primary antibody incubation, membranes were extensively washed with TBS-T, followed by 1 h incubation with HRP-conjugated secondary antibodies. Finally, membranes were washed and incubated with chemiluminescence reagents (ECL; GE Life Sciences) and visualized on a FujiFilm LAS-4000 Image Analysis system.

### *In vitro* kinase assays

In total, 300 ng of purified recombinant GST-tagged rat ArfGAP1 proteins (F5 252-359 fragment: WT, S284A, T291A or T292A) were incubated with 95 ng of recombinant GST-tagged human LRRK2 (residues 970-2527: G2019S or D1994A; Invitrogen) in 5 µl 10x kinase buffer (Cell Signaling Technology) and 2 µl [^33^P]-γ-ATP (0.2 µCi/reaction) in a final volume of 15 µl by shaking at 30°C for 30 min. Where indicated, an increasing concentration of F5 GST-ArfGAP1 protein (WT or 3A mutant; 75, 300, 500 or 750 ng) was added with LRRK2 (G2019S or D1994A) in the samples and incubated for 30 min in similar fashion. The reaction was terminated by adding 5 µl of 4x LDS sample buffer (Invitrogen) followed by denaturing the samples at 70°C for 10 min. Samples were resolved by SDS-PAGE, transferred onto PVDF membranes (GE Life Sciences) and subjected to autoradiographic detection using a Typhoon Phosphorimager (Cytiva) as described previously [74]. Membranes were subsequently blocked with TBS-T containing 5% milk and incubated with anti-GST-HRP antibody (GE Life Sciences). Proteins were visualized using enhanced chemiluminescence (GE Life Sciences) and a luminescent image analyzer (LAS-4000, FujiFilm). Similar kinase assays were conducted using recombinant full-length GST-ArfGAP1 and GST-ΔN-LRRK2 (WT, G2019S or D1994A) in the presence of excess cold ATP, for analysis by mass spectrometry. In separate radioactive kinase assays with [^33^P]-γ-ATP, we incubated recombinant GST-tagged ArfGAP1 deletion mutants (WT or F1-F5) with purified full-length FLAG-tagged LRRK2 (WT, G2019S or D1994A) derived by anti-FLAG IP from transfected SH-SY5Y cells, as described [75]. Membranes were first imaged for ^33^P by autoradiography using a Typhoon Phosphorimager, and then by Western blotting with anti-FLAG-HRP and anti-GST-HRP antibodies.

### Cortical neurite length assay

Rat primary cortical cultures were co-transfected at DIV 3 with FLAG-LRRK2 and DsRed-Max-N1 at a 10:1 molar ratio using Lipofectamine 2000 reagent (Invitrogen) according to manufacturer’s recommendations. For co-expression experiments, transfections were performed with FLAG-LRRK2, YFP-ArfGAP1 (WT, phospho-null / phospho-mimic mutants) and DsRed plasmids at a 10:10:1 molar ratio. At DIV 6, cultures were fixed with 4% paraformaldehyde and processed for immunocytochemistry with mouse anti-FLAG (M2) antibody (Sigma-Aldrich) and anti-mouse IgG-AlexaFluor-488 or -633 antibodies (Invitrogen). Fluorescent images were acquired, processed and analyzed for neurite length measurement similar to our previous study [16].

### Golgi fragmentation assay

SH-SY5Y cells were transiently transfected with YFP-ArfGAP1 (WT, phospho-null / phospho-mimic mutants), fixed and processed for immunocytochemistry with mouse anti-GM130 antibody and anti-mouse-IgG AlexaFluor-546. Golgi morphology was assessed by classifying complexes as either normal intact, partially fragmented, or fully fragmented, as previously reported [16].

### Mass spectrometry

We performed similar *in vitro* kinase assays with excess non-radioactive ATP and measured the phosphorylation of recombinant GST-tagged WT ArfGAP1 by GST-ΔN-LRRK2 (WT, G2019S and D1994A) using mass spectrometry. An in-solution digestion protocol was performed as detailed previously [76]. Briefly, samples were lysed, and proteins were reduced with dithiothreitol (DTT) followed by alkylation with iodoacetamide (IAA) in the dark. Protein digestion was performed with Lys-C (enzyme:substrate ratio of 1:100) (Wako) at room temperature for 3 h followed by an additional overnight digestion with Lys-C. The digestion was stopped by acidification and the mixture was desalted on reversed phase C18 StageTips prior to liquid chromatography tandem mass spectrometry (LC-MS/MS, Thermo Scientific Q-Exactive HF-X; Michigan State University Proteomics Facility, East Lansing, MI). In separate experiments, LC-MS/MS analysis was conducted to detect ArfGAP1 protein interactors in anti-GFP IP samples derived from SH-SY5Y cells transiently expressing YFP-tagged WT ArfGAP1 versus an empty plasmid control.

### Raw data processing

The mass spectrometric raw data were processed and analyzed using MaxQuant software (version 1.4.7.2) [77]. Proteins were detected using the implemented Andromeda search engine and human / rat database with common contaminants. Oxidation of methionine and acetylation of the protein N-terminus were selected for variable modifications, whereas carbamidomethylation of cysteines were chosen for fixed modification. Lys-C was taken as the protease where a maximum of 2 missed cleavages were allowed. For fragment ions, 0.5 Da mass tolerances were selected and for identification, only peptides with a minimum of six amino acids were considered. The false discovery rate was set to 1% on the peptide-spectrum-match and protein level using the implemented decoy algorithm. The minimum ratio count was set to 2. Data analysis and visualization were performed using Perseus software (Max Plank Institute, Martinsried) and the statistical environment R. Significant differentially-expressed phosphoproteins or proteins were identified by a permutation-based FDR approach using a cutoff of 0.01 and 500 permutations. Moreover, significant differentially expressed interacting proteins were identified by taking log2 fold change ≥ 2. Interaction networks, GO, KEGG and INTERPRO domain analysis were done by STRING [78], the statistical environment R and Perseus [79].

### Statistical analysis

All data were analyzed by two-tailed, unpaired Student’s *t*-test for pair-wise comparisons, one-way ANOVA with Dunnett’s multiple comparisons test for samples grouped by one factor, or two-way ANOVA with Tukey’s multiple comparisons test for samples grouped by two factors, as indicated. *P*<0.05 was considered significant.

## Acknowledgements

This work was supported by a grant from the National Institutes of Health (NIH) R01NS091719 to D.J.M. We thank Amelie Moore for assistance with graphical illustration.

## Supplementary Data

**Figure S1. Impact of single phosphorylation mutants on ArfGAP1-induced Golgi fragmentation in neural cells. A**) SH-SY5Y cells transiently expressing YFP-tagged WT ArfGAP1 or empty vector were fixed and subjected to immunofluorescence with antibodies to different Golgi membrane markers (GM130, TGN46, Giantin, GOLGA4). Quantitation of Golgi morphology in ArfGAP1-positive cells reveals similar levels of Golgi fragmentation between Golgi markers (90-100% cells). **B-C**) Quantitation of Golgi fragmentation induced by overexpression of WT or single phospho-mutants of YFP-ArfGAP1 in SH-SY5Y cells. Single phospho-null or phospho-mimic mutants induce similar Golgi fragmentation levels to WT ArfGAP1 (∼85% cells). **D**) Golgi fragmentation induced by ArfGAP1 double (T291A/T292A) or triple (3A) phospho-null mutants indicates partial Golgi fragmentation of double mutants (∼65% cells) or 3A mutant (∼45% cells) compared to WT ArfGAP1 (∼85% cells). In each graph, bars represent the percentage of cells with normal or fragmented Golgi from a representative experiment.

**Figure S2. Impact of single phospho-null mutants on ArfGAP1-induced inhibition of neurite outgrowth. A**) Rat primary cortical neurons were co-transfected at DIV3 with YFP-ArfGAP1 (WT, S284A, T291A or T292A) or empty vector and DsRed-Max-N1 plasmids, and fixed at DIV6 for confocal fluorescence microscopy analysis. Fluorescent images reveal the co-labeling of cortical neurons with YFP-ArfGAP1 (green) and DsRed (red), with the DsRed images pseudo colored (ICA) to enhance the contrast of neuritic processes. Neuronal soma (white arrows) and axonal processes (black arrowheads) are indicated. Scale bars: 50 μm. **B**) Quantitative analysis of DsRed-positive axon length in YFP-ArfGAP1-positive neurons or control neurons (empty vector) is shown. Bars represent the mean ± SEM axon length (in μm) from 90-120 double DsRed-/YFP-ArfGAP1-positive neurons, or single DsRed-positive neurons (control), across at least three independent experiments/cultures. \*\*\*\**P*<0.0001 compared to control (DsRed alone) by one-way ANOVA with Dunnett’s multiple comparisons test. *ns*, non-significant.

**Figure S3. Impact of single phospho-mimic mutants on ArfGAP1-induced inhibition of neurite outgrowth. A**) Rat primary cortical neurons were co-transfected at DIV3 with YFP-ArfGAP1 (WT, S284D, T291D or T292D) or empty vector and DsRed-Max-N1 plasmids, and fixed at DIV6 for confocal fluorescence microscopy analysis. Fluorescent images reveal the co-labeling of cortical neurons with YFP-ArfGAP1 (green) and DsRed (red), with the DsRed images pseudo colored (ICA) to enhance the contrast of neuritic processes. Neuronal soma (white arrows) and axonal processes (black arrowheads) are indicated. Scale bars: 50 μm. **B**) Quantitative analysis of DsRed-positive axon length in YFP-ArfGAP1-positive neurons or control neurons (empty vector) is shown. Bars represent the mean ± SEM axon length (in μm) from 90-120 double DsRed-/YFP-ArfGAP1-positive neurons, or single DsRed-positive neurons (control), across at least three independent experiments/cultures. \*\*\**P*<0.001 compared to control (DsRed alone) by one-way ANOVA with Dunnett’s multiple comparisons test. *ns*, non-significant.

**Figure S4. Effects of ArfGAP1 or LRRK2 expression alone on neurite outgrowth. A**) Rat primary cortical neurons were co-transfected at DIV3 with YFP-ArfGAP1 (WT), FLAG-LRRK2 (WT or G2019S) or empty vector and DsRed-Max-N1 plasmids, and fixed at DIV6 for immunofluorescence with anti-FLAG antibody. Fluorescent images reveal the co-labeling of cortical neurons with YFP-ArfGAP1 (green) or FLAG-LRRK2 (yellow) and DsRed (red), with the DsRed images pseudo colored (ICA) to enhance the contrast of neuritic processes. Neuronal soma (white arrows) and axonal processes (black arrowheads) are indicated. Scale bars: 50 μm. **B**) Quantitative analysis of DsRed-positive axon length is shown in YFP-ArfGAP1-positive, FLAG-LRRK2-positive or control (empty vector) neurons. Bars represent the mean ± SEM axon length (in μm) from 90-120 double DsRed-/ArfGAP1-positive or DsRed-/LRRK2-positive neurons, or single DsRed-positive neurons (control), across at least three independent experiments/cultures. \*\*\*\**P*<0.0001 compared to control (DsRed alone) by one-way ANOVA with Dunnett’s multiple comparisons test. *ns*, non-significant.

