## Supplementary figures and images for "LRRK2 regulates ArfGAP1 membrane localization, activity and neuronal toxicity via phosphorylation within its lipid-sensing ALPS2 motif"

### Figure S1

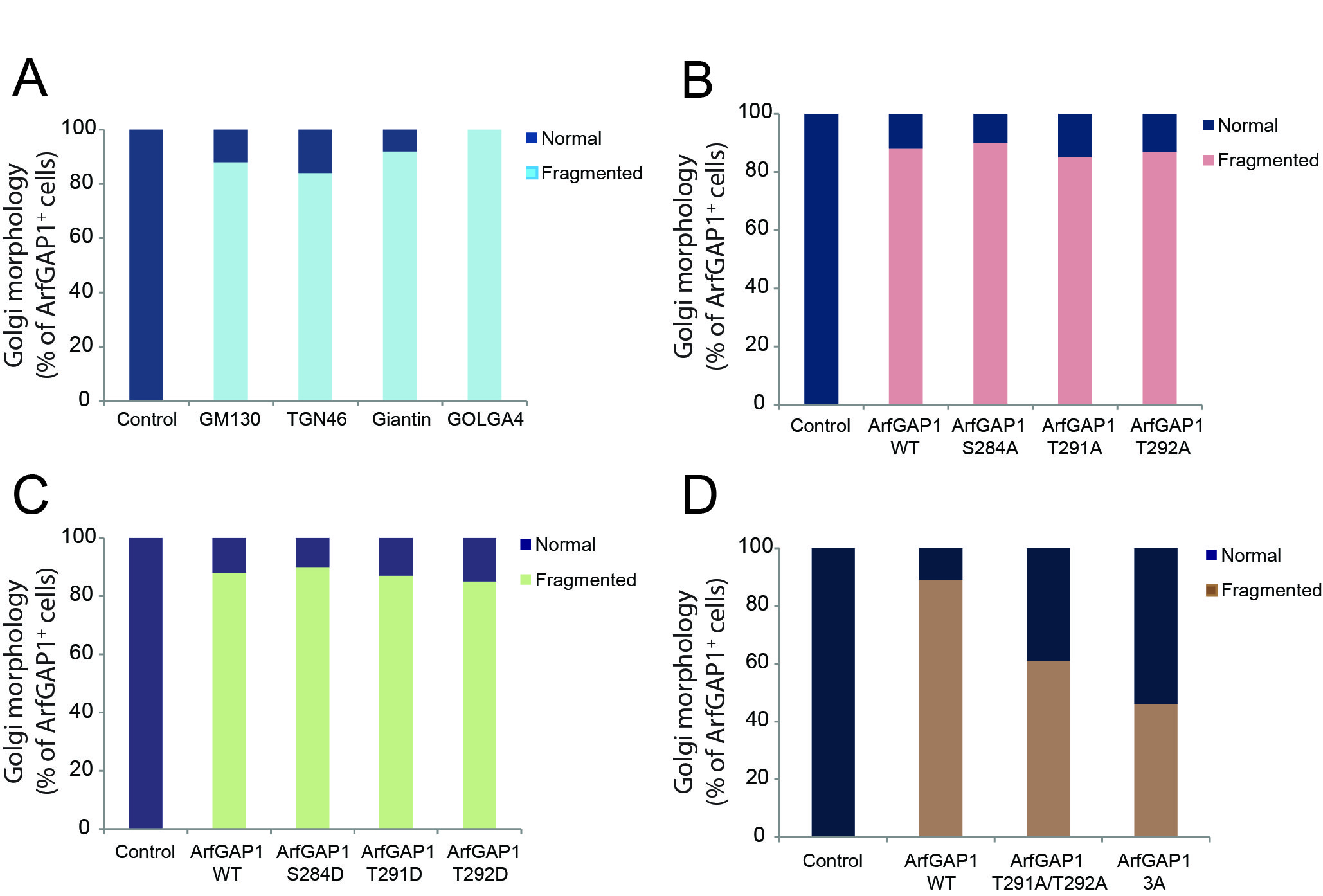

### Figure S2

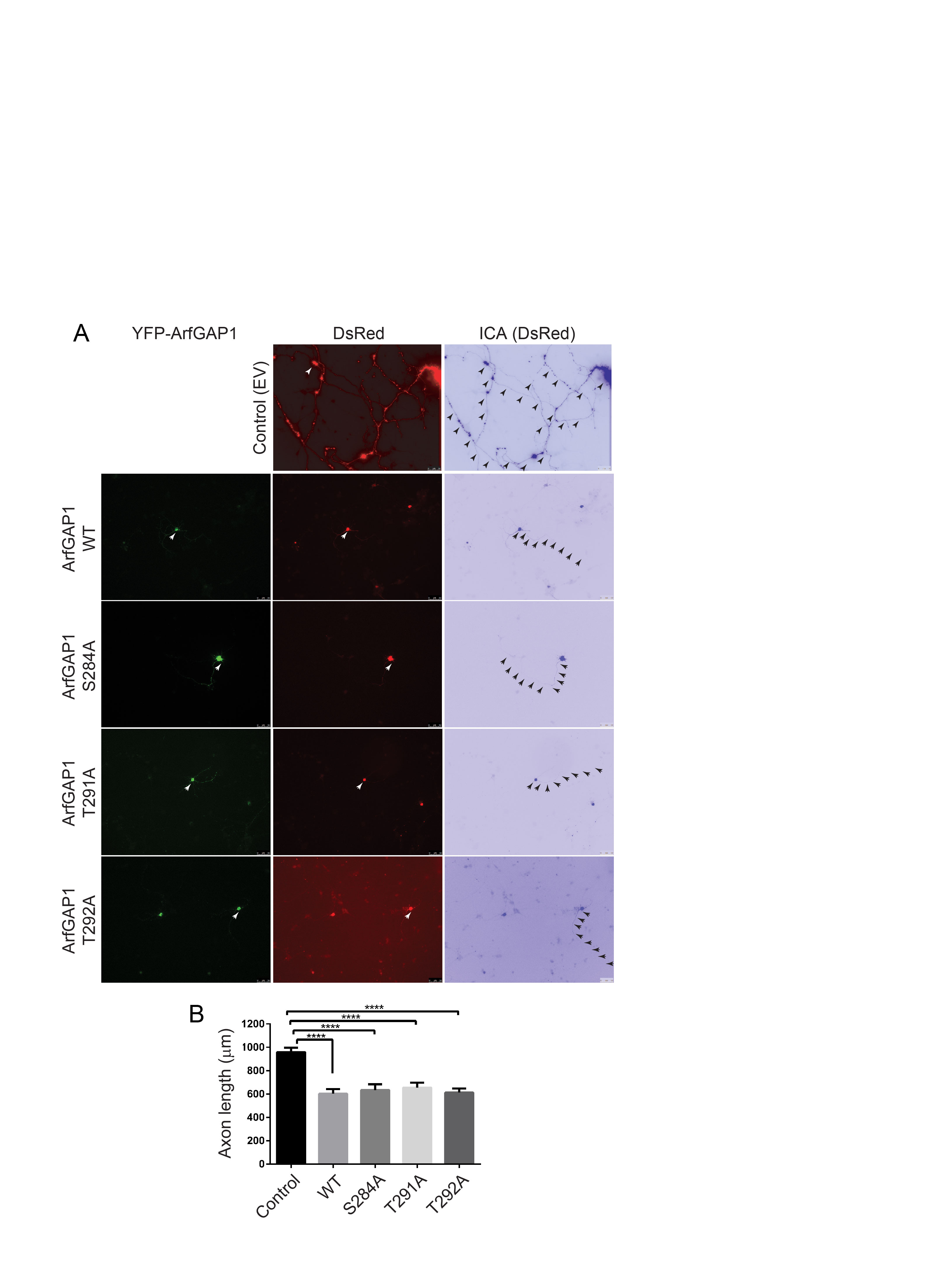

### Figure S3

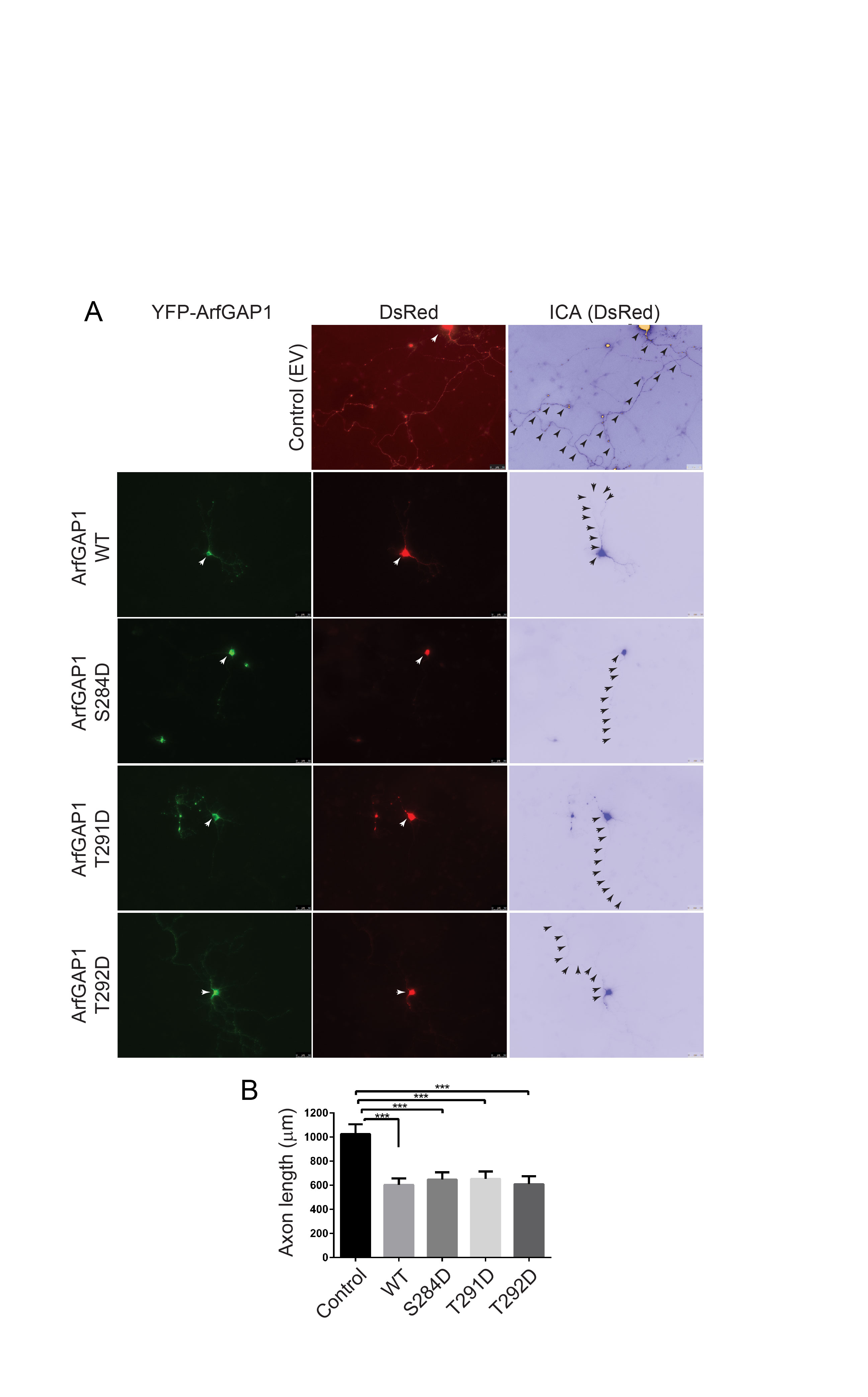

### Figure S4

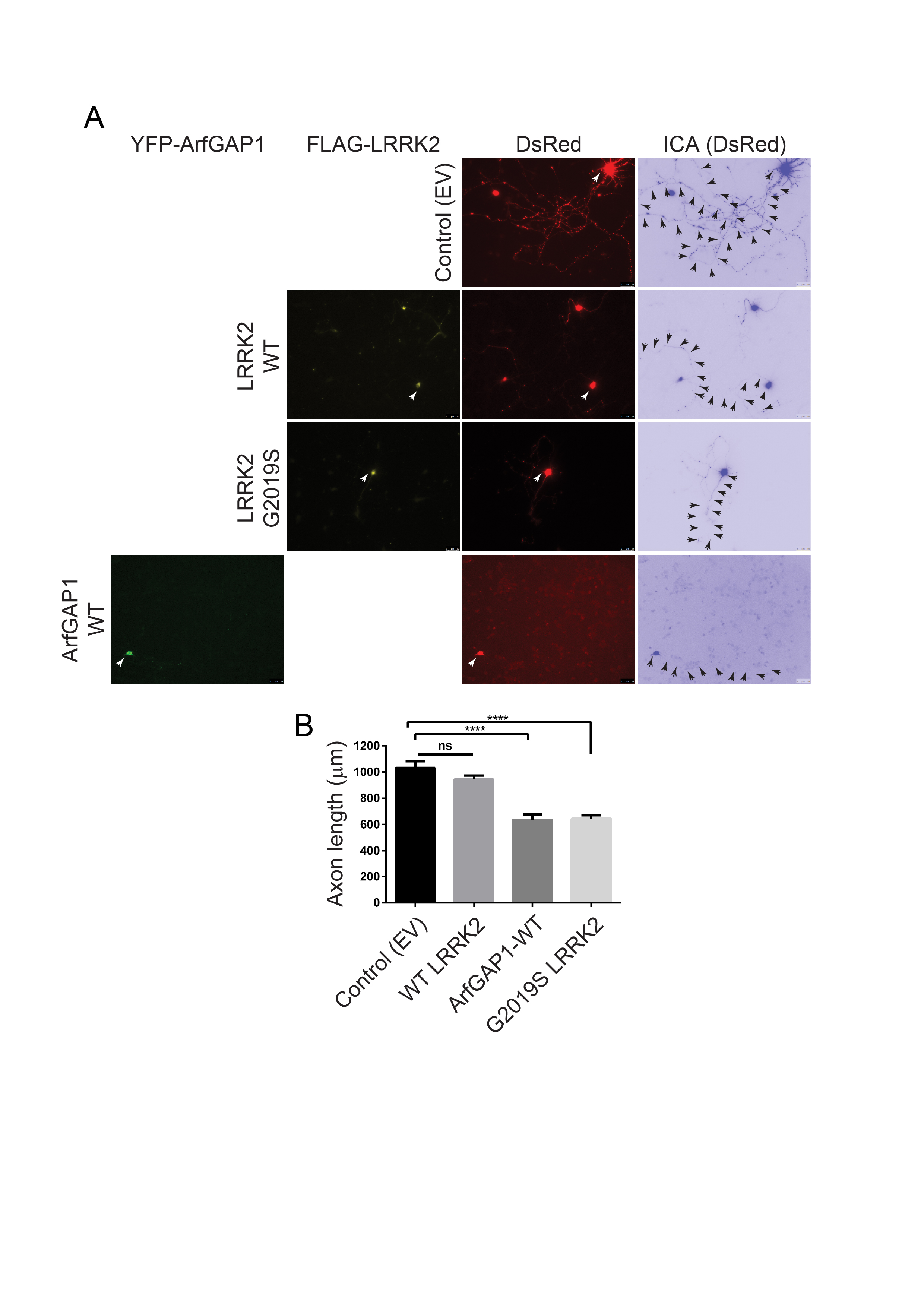
